# Encoding of wind direction by central neurons in *Drosophila*

**DOI:** 10.1101/504753

**Authors:** Marie P. Suver, Andrew M. M. Matheson, Sinekdha Sarkar, Matthew Damiata, David Schoppik, Katherine I. Nagel

## Abstract

Wind is a major navigational cue for insects, but how wind direction is decoded by central neurons in the insect brain is unknown. Here, we find that walking flies combine signals from both antennae to orient to wind during olfactory search behavior. Movements of single antennae are ambiguous with respect to wind direction, but the difference between left and right antennal displacements yields a linear code for wind direction in azimuth. Second-order mechanosensory neurons share the ambiguous responses of single antenna and receive input primarily from the ipsilateral antenna. Finally, we identify a novel set of neurons, which we call wedge projection neurons, that integrate signals across the two antennae and receive input from at least three classes of second-order neurons to produce a more linear representation of wind direction. This study establishes how a feature of the sensory environment – the wind direction – is decoded by single neurons that compare information across two sensors.

## INTRODUCTION

Animals have evolved a broad array of systems that enable them to extract relevant features of their sensory environment. These systems often achieve accurate encoding of the sensory world by comparing signals between sensors. For instance, in the auditory system of birds and mammals, coincidence detectors compare activity between neural ‘delay lines’ that receive inputs from opposite side of the head to compute sound direction (Konishi, 2003). Alternatively, perception can be enabled by comparing relative activities across sensors in the same region of space, but with different tuning properties. This occurs in touch perception in mammals, where information from mechanosensors with differing tuning properties in the same region of skin is integrated in the spinal cord (Abraira & Ginty, 2013). However, the central processing of sensory information is often challenging to study in vertebrates, particularly with single-cell precision.

One of the most iconic examples of paired sensors in the animal kingdom are arthropod antennae. Insects use their antennae for olfaction and thermosensation, and to perform a wide variety of mechanosensory behaviors. For instance, cockroaches use mechanosensory information from their antenna to guide high-speed locomotor activity (Camhi & Johnson, 1999), and to actively probe objects (Okada & Toh, 2006). Many insects also use their antennae to sense wind direction, an important cue for navigation. Examples include cockroaches, which can use wind to set a straight course in the absence of visual landmarks (Bell & Kramer, 1979), and desert ants that use wind cues to navigate towards a learned food source (Müller & Wehner, 2007). The ability to gauge wind direction is critical in many species for navigating towards an odor source (Wolf and Wehner 2000, Willis and Avondet, 2005, Bell and Wilson, 2016) because wind often provides a stronger direction cue than odor concentration in a turbulent environment (Murlis, Willis, Carde, 2000; Carde and Willis, 2008). Orientation to wind is abolished by stabilizing the antennae in cockroaches (Bell & Kramer, 1979), ants (Wolf & Wehner, 2000), and walking *Drosophila* (Álvarez-Salvado et al., 2018), indicating a critical role for antennal mechanoreceptors. Furthermore, experiments in flying flies suggest that information from both antennae must be combined to accurately gauge wind direction (Budick, Reiser, & Dickinson, 2007). This analysis is supported by video recordings of antennal movements in response to wind, which suggest that the two antennae are differentially displaced depending on the wind direction (Patella & Wilson, 2018; Yorozu et al., 2009). However, the neural circuits underlying this computation are not known.

In insects, movements of the antennae are detected by multiple sensors, including a large set of stretch receptors located in the second antennal segment, collectively known as the ‘Johnston’s Organ’ (JO; Johnston, 1855). Movements of the third antennal segment relative to the second are aided by a feather-like structure called the arista that acts as a sail and enables the fly to detect different types of mechanical stimuli including conspecific song (Göpfert & Robert, 2002). About 500 primary mechanosensory neurons, known as ‘Johnston’s Organ neurons’ (JONs) project from the antennae through the antennal nerve towards the central brain, to a region called the antennal mechanosensory and motor center (AMMC; Kamikouchi, Shimada and Ito, 2006). Traditionally, JONs have been broadly grouped into phasic (AB JONS) and tonic (CE JONs) classes that project to different regions of the AMMC and carry auditory (AB) or wind and gravity (CE) information (Kamikouchi et al., 2009; Yorozu et al., 2009). However, D JONs carry both phasic and tonic information (Matsuo et al. 2014), and it is likely that these broad classes are composed of many more specialized cell types (Kamikouchi et al., 2006; Patella & Wilson, 2018).

In the AMMC, mechanosensory information is processed by multiple second-order neurons, many of which project to the nearby ‘wedge’ (WED), a major third-order mechanosensory center (Kamikouchi et al., 2009; Lai, Lo, Dickson, & Chiang, 2012; Matsuo et al., 2016a; Patella & Wilson, 2018; Vaughan, Zhou, Manoli, & Baker, 2014). Among these second-order neurons, two classes known as APN2 and APN3 are thought to lie anatomically downstream of wind-sensitive JONs (Matsuo et al., 2016a; Vaughan et al., 2014). APN3 responds to tonic displacements of the antenna induced by a piezo probe (Chang et al., 2016), while APN2 has not been physiologically characterized. In contrast, the contralaterally projecting B1 neurons (also called APN1 in Vaughan *et al*., 2014) are thought to be downstream of the phasic AB JONs (Chang et al., 2016; Lai et al., 2012). B1 neurons were functionally implicated in the perception of courtship song in a previous study, while APN2 and 3 were found to be dispensable for this behavior (Vaughan et al., 2014). This suggests that the functional segregation of phasic auditory and tonic wind signals is maintained in second-order neurons. However, APN3 neurons respond to tonic displacements of the antenna as well as sound and sound-like stimuli (Chang et al., 2016). Furthermore, B1 neurons exhibit directionally tuned responses to piezo displacements (Azevedo & Wilson, 2017), complicating this simple picture. The responses of these neurons to wind, as opposed to piezo displacements of a single antenna, have not been measured.

Where might wind direction be computed from information about the displacements of the two antennae? A recent pan-neuronal imaging study in *Drosophila* suggested that information from the two antennae is first combined in the WED (Patella & Wilson, 2018). Because this study used pan-neuronal drivers to measure activity, however, it was not able to establish the identity of any specific neurons involved in this process or identify higher-order areas involved in wind encoding. In the central brain, neurons responding to wind stimuli and to tonic deflections of the antennae have been described in the central complex (in bees: Homberg, 1985, locusts: Homberg, 1994, and cockroaches: Ritzmann, Ridgel and Pollack, 2008). However, there is a gap in our understanding of how information reaches these regions of the brain from peripheral processing centers in the AMMC and WED.

In this study, we use whole cell electrophysiology to identify specific WED projection neurons that compute wind direction from displacements of the two antennae and to elucidate presynaptic partners underlying this computation. We first show that walking flies combine information from both antennae to effectively orient to wind. Next, we perform a quantitative analysis of antennal movements in response to directional wind and show that each antenna provides an ambiguous signal about wind direction, but that this ambiguity can be resolved by a simple linear combination of displacement signals from the two antennae. Whole cell recordings demonstrate that second order neurons encode displacements of the ipsilateral antenna with a similar ambiguity regarding wind direction as single antenna. Then, we perform a whole-cell electrophysiology screen and identify novel wedge projection neurons (WPNs) that integrate information across the two antennae to generate a more linear encoding of wind direction in azimuth. Finally, we use pharmacological and genetic silencing experiments, along with trans-synaptic labeling, to show that multiple populations of second order neurons provide input to WPNs, and that these overlap with populations previously shown to play a role in auditory encoding and courtship behavior. In this study, we introduce novel neurons in the insect brain that integrate signals across the antennae to encode wind direction, identify new regions of the dorsal brain as putative centers of mechanosensory processing, and provide insight into the circuit mechanisms by which mechanosensory neurons compute important features of the mechanosensory environment. Our results illustrate circuit mechanisms for decoding a salient feature of the environment – wind direction – from the activities of paired sensors.

## RESULTS

### Robust wind orientation behavior requires both antennae

Walking *Drosophila* rely on antennal movements both to orient upwind in response to odor, and downwind in the absence of odor (Álvarez-Salvado et al., 2018). However, it is not known whether information from both antennae is required for these behaviors. To study how the two antennae work together to promote orientation to wind, we measured the behavioral responses of freely walking flies to wind and stabilized either one or both antennae (Figure 1A). As shown previously (Álvarez-Salvado et al., 2018), flies orient upwind in the presence of odor, and this orientation is blocked by stabilizing both antennae (Figures 1B-D). In contrast, local search behavior produced at odor offset is not impaired by antenna stabilization, indicating that flies with stabilized antennae can still detect the odor (Figures 1B, 1E, and 1F). In the absence of odor (blank trials), flies tend to aggregate at the downwind end of the tunnel; this downwind preference is lost when antennal inputs are blocked (Figure 1G).

**Figure 1.**
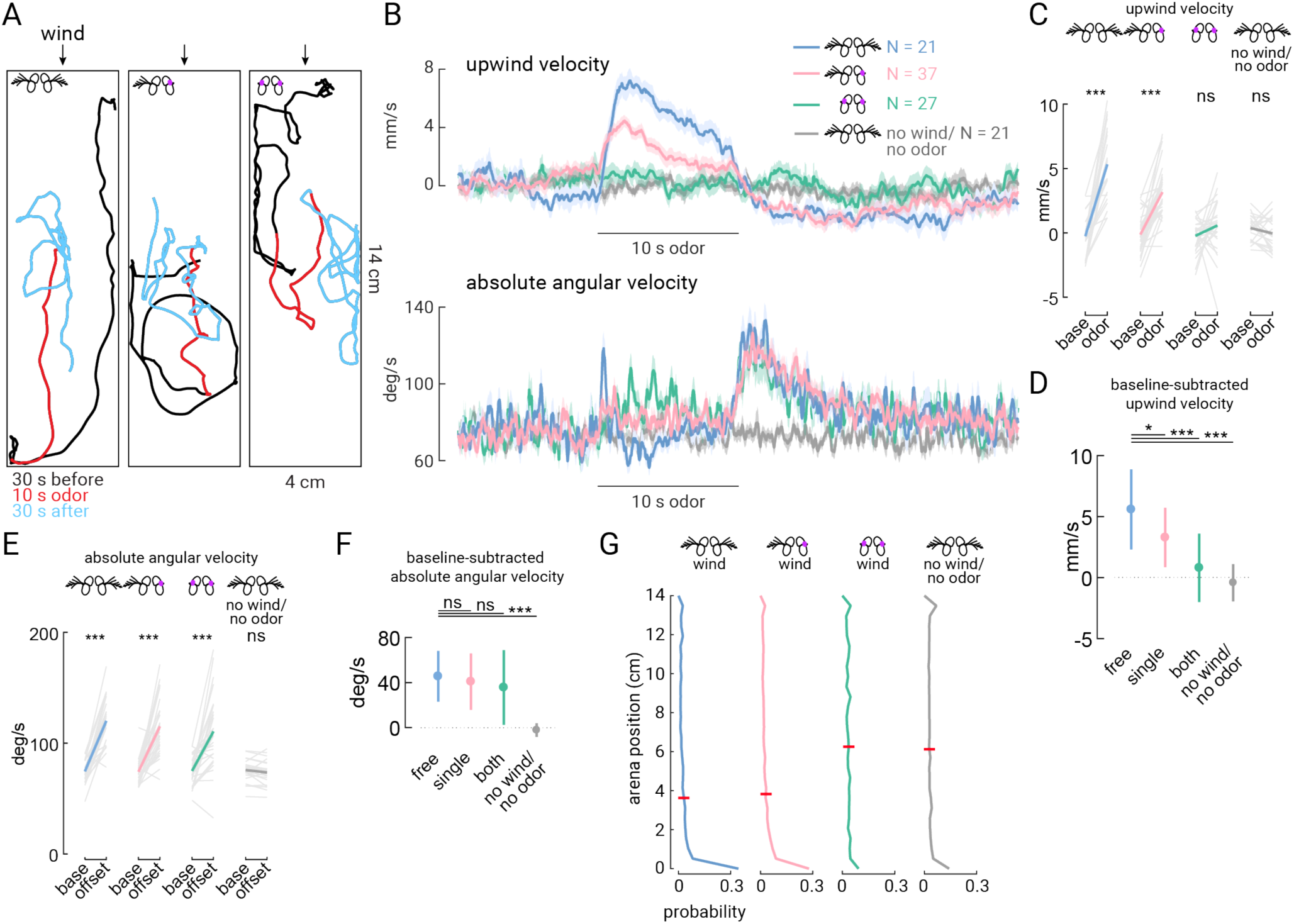
Both antennae contribute to odor-driven upwind orientation. (A) Example walking trajectories during single trials consisting of wind only (black, 30 s), wind with odor (1% apple cider vinegar, red, 10 s), and wind only again (cyan, 30 s). Wind was delivered from the top of the arena (top of this image). Examples are shown for flies with intact antennae (left), one antenna stabilized (center) or both antennae stabilized (right). (B) Top: Upwind velocity (average across flies +/− standard error) in response to a 10 s odor pulse. Responses from flies with intact antennae are shown in blue (N=22 flies), with one antenna stabilized in pink (N=37), both stabilized in green (N=27), and intact antenna but no wind and no odor; (N = 21). Bottom: absolute angular velocity (average across flies +/− standard error) for the same flies as above. (C) Odor-evoked upwind velocity for single flies (average of 5 s during odor versus 15 s pre-odor baseline). Flies with intact antennae or one antennae stabilized exhibit increased upwind velocity during the odor, but flies with both antennae stabilized or in the absense of wind and odor do not (Mann-Whitney U test, P = 0.000040, 0.00000067, 0.11, and 0.22, respectively). (D) Average baseline-subtracted upwind velocity responses +/− standard deviation for the single-fly averages plotted in (C). On average, flies with one or both antenna stabilized, and in the absense of wind and odor, walk upwind less than flies with free antennae (Wilcoxon signed-rank test, P = 0.016, 0.0000012, and 0.000000082, respectively). In the absense of wind, upwind velocity is less than (-0.60 mm/s), but not significantly different than zero (Wilcoxon signed-rank test, P = 0.22). (E) Odor-offset evoked increase in absolute angular velocity for single flies (average of 2 s post-odor versus 15 s pre-odor baseline). Flies in all treatments in the presense of wind exhibit an increase in angular speed after odor offset, but not in the absense of wind (Mann-Whitney U test, P = 0.000053, 0.00000017, 0.000026, and 0.26). (F) Average baseline-subtracted absolute angular velocity +/− standard deviation across the single flies plotted in (E). Flies did not exhibit any significant change in angular speed with stabilized antennae (Wilcoxon signed-rank test, P = 0.33, 0.18). In the absense of wind, flies do not exhibit any significant chance in baseline-subtracted angular velocity (Wilcoxon signed-rank test, P = 0.0262). (G) Probability distribution of flies along the length of the walking arena during the wind-only stimulus (no odor) for flies with free antenna (N=22 flies), free antenna but no wind and no odor (N = 21), single antenna stabilized (N=36), and both antenna stabilized (N=26) Wind comes from the top. Black distribution shows antenna-free fly responses in the arena with no wind (N=22). Red bars indicate average position. Mean position of flies is significantly different between no wind and wind (grey vs. blue, Mann-Whitney U test, P = 0.000034). Mean position of flies with one antenna stabilized is not significantly different from flies with free antenna during wind (pink vs. blue, P = 0.29). Mean position of flies with both antenna stabilized during wind is not different from the position of flies with intact antenna and no wind (green vs. grey, P = 0.93). All comparisons were corrected with the Bonferroni method.

To examine whether input from both antennae is required for wind orientation, we stabilized either the left or right antenna and again measured flies’ responses to odor pulses. We were unable to distinguish between flies walking on the ceiling or the floor in our arena, so we combined data from flies with right or left antenna stabilized (which were not statistically different; data not shown). Stabilization of a single antenna reduced upwind velocity during odor by about half on average (Figure 1D). Local search after odor offset was unaffected, similar to flies that had both antennae glued (Figures 1E and 1F). Flies with one antenna glued also exhibited no change in average downwind aggregation (Figure 1G), perhaps because this measure reflected integrated downwind orientation over time. Together these findings suggest that flies require both antennae to perform normal upwind orientation in response to odor, but that a single antenna is sufficient to produce the baseline downwind orientation. These results led us to ask how wind direction signals are encoded by movements of a single antenna compared with combined antennae movements.

### Single antenna motions are ambiguous with respect to wind direction

A previous study (Yorozu et al., 2009) showed that movements of the two antennae together can encode wind direction: wind from the front pushes both antennae towards the head, while wind from one side pulls the ipsilateral antenna away, and pushes the contralateral antenna towards the head. To examine how antennal displacements encode wind direction more quantitatively, we designed a stimulus apparatus to deliver wind to the fly from 5 different directions (Figures 2A and 2B). Importantly, the fly was mounted in a holder with a fixed position relative to the stimulus ports, ensuring reliable positioning of the fly at the center of the five wind streams across experiments. We verified that the velocity of the wind was identical across directions, so that any difference in antennal movements would be due to directionality rather than velocity (Figure 2C).

**Figure 2.**
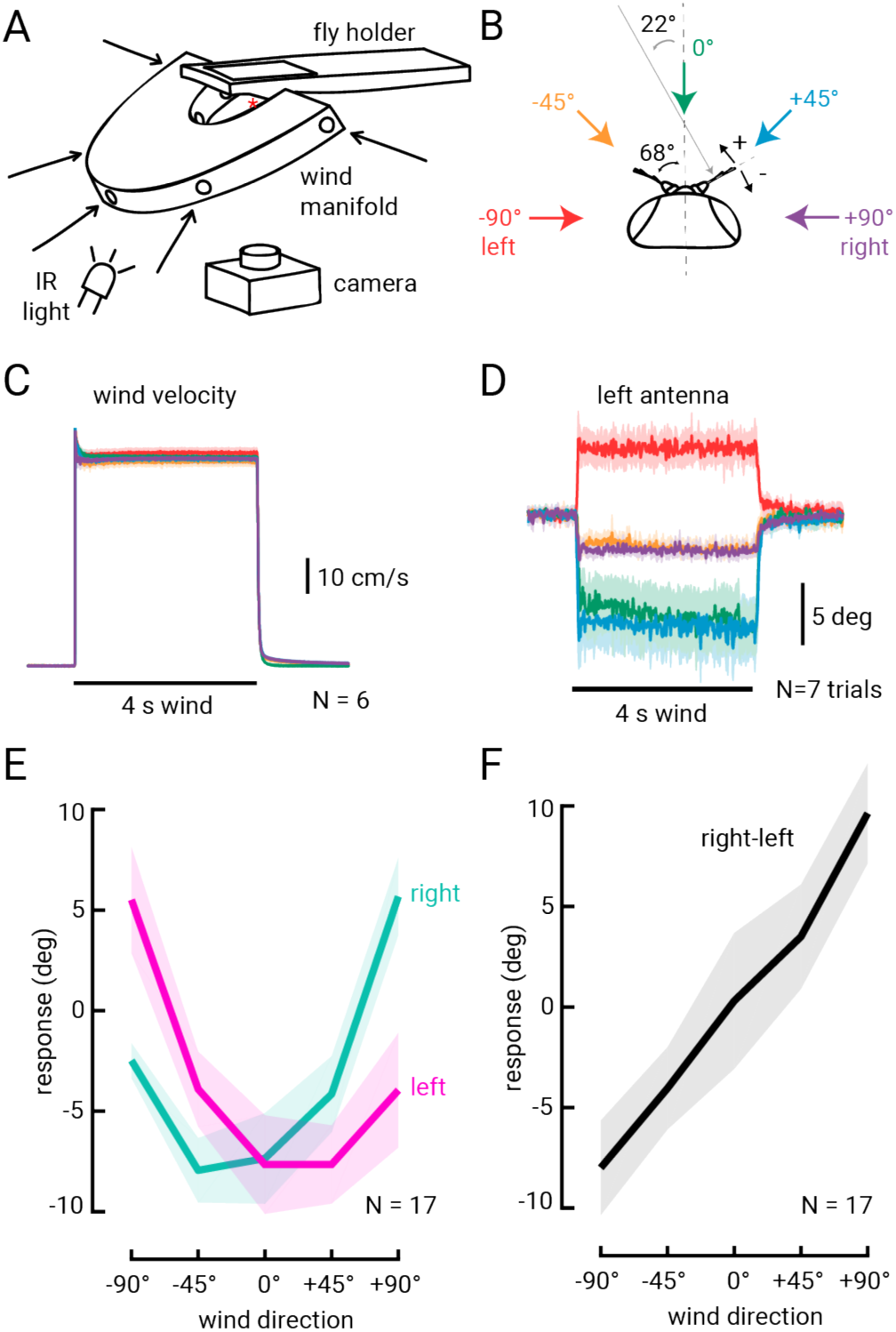
Wind direction encoding by antennal displacement. (A) Schematic of the wind stimulus apparatus. Wind is delivered to the fly from one of five directions through a manifold. The fly is mounted in a holder so that its head is centered at the intersection of the five channels. A camera is mounted below the fly, and the fly is lit by an infrared (IR) light aimed towards the fly from below. (B) Schematic of the fly head and antennae. We deliver wind to the fly from −90° (left), −45°, 0°, +45° and +90° (right) relative to the midline of the fly (dashed vertical line). We track antennal motions by measuring deflections of the aristae relative to the midline of the fly. Average resting inter-arista angle is 136° +/− 15°, and the wind angle normal to the arista’s axis is noted (22°) and indicated by the grey arrow. (C) Average wind velocity measured by a hot wire anemometer placed at the location of the fly head during experiments. Average steady state windspeed is 60 +/− 1.4 cm/s. Standard deviation across N=6 measurements is plotted as a shaded region behind the averages (but is difficult to discern because they lie on top of one another). (D) Average left arista deflections across N=7 trials for each of the 5 wind directions for one fly. (E) Average steady state deflections (+/− standard deviation) relative to baseline position for the left (pink) and right (teal) antennae across N=17 flies. Averages are derived from 5 frames (83 m s total) directly before and at the end of the wind stimulus (see Methods). (F) Difference in antennal deflections between the two antennae (right-left, from the same flies as plotted in (E)) is plotted in black. Standard deviation of the mean is plotted as a lighter shade around left, right, and right-left average deflections in (E) and (F).

We presented a 4 s tonic wind stimulus in random order with respect to direction, and recorded movements of the antenna using a camera mounted directly below the fly. By tracking the antenna’s position relative to the midline of the fly (Figure 2B), we found that the antennae are tonically displaced by the wind stimulus, remain deflected for the duration of the stimulus, then quickly relax back to their pre-wind position at stimulus offset (Figure 2D). We next examined displacements of the antennae as a function of wind orientation (Figures 2E and 2F). Each antenna exhibits a curved displacement function with the greatest displacement towards the head (minimum of the curve) at a point between 0° and 45° contralateral to that antenna. The right arista is positioned at approximately 68° relative to the midline at rest (Figure 2B); thus, displacement of the antenna towards the head should be maximal at ~−22° (normal to the arista), and displacement should decrease as the cosine of the angle as wind deviates from this direction. We observe exactly this relationship in the displacements of a single antenna (Figure 2E), each of which has a hook-shaped tuning curve for wind direction. However, because the two aristae are positioned at ~135° to one another, a nearly linear encoding of wind direction can be obtained by simply taking the difference between the two antennal displacements, as illustrated in Figure 2F. This analysis suggests that a single antenna provides an ambiguous representation of wind direction in azimuth, whereas a difference between the two antennae provides a more linear encoding of wind direction. The difference between antennal displacements also provides greater resolution for wind angles directly in front of the fly, where a single antenna provides a weak and ambiguous signal. These results raise the question of whether there are neurons in the brain that generate such representations.

### Two classes of AMMC projection neurons encode tonic displacements of the ipsilateral antenna

We next sought to characterize how antennal movements are represented by neurons in the central brain to encode wind direction. The AMMC projection neurons APN2 and APN3 are thought to be downstream of the wind-sensitive CE JONs (Chang et al., 2016; Vaughan et al., 2014), so we investigated the tuning of these neurons first. We performed whole cell patch clamp recordings from single, genetically identified APN2 or APN3 neurons from both the left or right side of the brain. Responses from neurons on the left side of the brain were flipped and averaged with neurons from the right side, and we describe neural responses to wind as ipsilateral or contralateral relative to cell body location. We presented 4 s tonic wind stimuli from five directions using the same apparatus described above (also see Figure 2). We found that APN2 neurons were tonically inhibited by wind from all directions except the ipsilateral side (Figures 3B-D). APN2 neurons were non-spiking and responded to tonic wind stimulation with graded changes in membrane potential and little adaptation (Figures 3B-D). All APN2 cells we recorded were least inhibited by ipsilateral wind, although we observed some variation in the amount of inhibition across cells (Figure 3D). The tuning curves for APN2 neurons had a hooked shape (Figure 3D), much like the tuning curve for ipsilateral antennal displacement.

**Figure 3.**
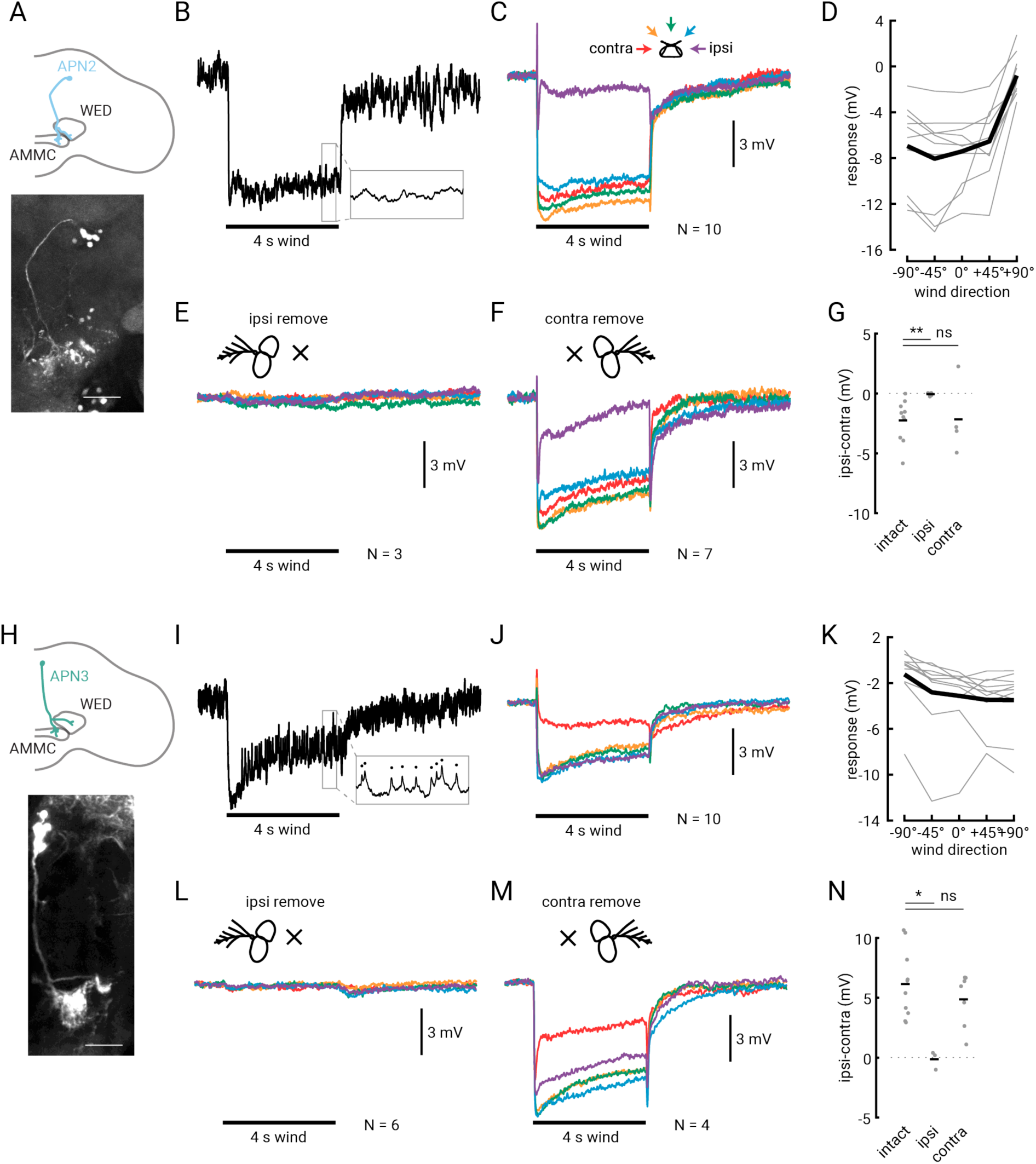
Second-order APNs encode tonic deflections of the ipsilateral antenna. (A) Schematic of APN2 anatomy (blue) on one half of the brain (gray outline), with processes in the AMMC and the WED. Below: GFP signal in *24C06-GAL4*, the line we used to target APN2, in the right hemisphere of the brain. Scale bar is 30 μΜ. (B) Single raw trace of an intracellular recording from APN2 in response to a 4 s wind stimulus (black bar) delivered from 0°. Inset shows a 50 ms expanded trace. (C) Average membrane potential response across N=10 APN2 neurons (from 8 flies) to the five wind directions (color convention shown above the traces). (D) Average steady-state tuning (last 1 s of wind stimulus relative to last 1 s of baseline) for single cells (thin gray lines) and across flies (thick black line). (E) Average APN2 response to wind from five directions after ipsilateral antenna removal. (F) Average APN2 response to wind after contralateral antenna removal. (G) Response range (ipsilateral-contralateral wind response) for flies with intact antennae or with the ipsilateral or contralateral antenna removed. Single-fly differences plotted as grey dots; black bars indicate average across flies. Antennae-intact and ipsilateral removed are significantly different (two-sample student’s t-test, P = 0.0037). Antennae-intact and contralateral removed are not significantly different (two-sample t-test, P = 0.34). (H) Schematic of APN3 (green) depicted in the right hemisphere, with processes in the AMMC and WED. Below: GFP expression in *70G01-GAL4*, the line we used to target APN3. Scale bar is 30 μΜ. **i**, Single raw intracellular response of one APN3 to wind from +45°. Inset shows small spikes, indicated by the black dots, that occur in 50 ms segment during the wind stimulus. (J) Average membrane potential response of 10 APN3 neurons (from 10 different flies) to wind from the five directions. (K) Steady state tuning of APN3 for single cells (thin lines) and across flies (thick line), computed in the same way as in (D). (L) Average APN3 response to wind from five directions after ipsilateral antenna removal. (M) Average APN3 response to wind after contralateral antenna removal. (N) APN3 response range (ipsilateral-contralateral wind response) for flies with intact antennae or with the ipsilateral or contralateral antenna removed. Symbols as in (G). Antennae-intact and ipsilateral removed are significantly different from one another (two-sample student’s t-test, P = 0.0093). Antennae-intact and contralateral removed are not significantly different (two-sample t-test, P = 0.94). All response traces are plotted on the same scale, and number of cells (each in a different fly) is listed to the right of the stimulus bar.

In contrast, APN3 neurons were inhibited by wind from all directions except the contralateral side (Figures 3I-K, Figure S1), the opposite of the ipsilateral antenna’s wind tuning. APN3 neurons produced small-amplitude spikes (28.6 Hz baseline, Figure 3I) as reported previously. APN3 responses were generally smaller and more variable than those of APN2 neurons (Figures 3I-K); there are also more APN3 neurons labeled in the driver line we used to target them (approximately 11 per hemisphere in *70G01-GAL4*) than in the driver used to target APN2 (approximately 6 per hemisphere labeled by *24C06-GAL4*).

Given these results, we wondered whether APNs receive any input from the contralateral antenna, or whether they are solely ipsilateral. A recent study (Chang et al., 2016) found that APN3 did not respond to piezo displacements of the contralateral antenna. Thus, we removed either the ipsilateral or contralateral antenna, and then measured the responses of APN2 and APN3 to the five wind directions. When we removed the ipsilateral antenna, we found that this entirely abolished the wind response in both APN2 and APN3 (Figure 3E, 3G, 3L and 3N; two-sample t-test, P = 0.0037 and 0.011, relatively). Removal of the contralateral antenna, however, had no significant effect on the tuning or magnitude of the response in either neuron (Figure 3F, 3G, 3M, and 3N, P = 0.34 and 0.94, relatively). We did notice a slight increase in adaptation in APN2 neurons when the contralateral antenna was removed (Figure 3F), and a slight increase in response magnitude in APN3, however not all flies exhibited these changes (data not shown), so we think they may simply represent variability across cells. Given these results, we conclude that both of these populations of AMMC projection neurons encode displacements of the ipsilateral antenna, suggesting that integration across the two antennae occurs in downstream circuits.

### A class of wedge projection neurons integrates information from the two antennae to generate a linear encoding of wind direction

To identify wind-sensitive neurons downstream of the APNs that might integrate information from the two antennae, we performed an electrophysiological screen of projection neurons with putative input processes in or near the wedge (Jenett et al., 2012). We measured the wind responses of many candidate neurons (26 distinct classes), and found few that were both tonically activated by wind and exhibited strong directional tuning (Figure S2). However, we discovered one class of neurons, which we call ‘wedge projection neurons’ (WPNs), that consistently responded to wind stimuli in a tonic, directional manner (Figure 4). The line we found (*70B12-GAL4*) strongly labeled only two of these neurons on each side of the brain, and the two cells were anatomically and physiologically similar. By intracellularly filling two ipsilateral WPNs (Figures 4A and 4B), as well as staining for pre-synaptic markers (Figure S3), we found that these cells receive putative input in the wedge, and project dorsally to the posterior lateral protocerebrum (PLP), where one arm of the neuron branches off. The main branch then projects dorsomedially towards the superior clamp (SCL) and antler (ATL), before crossing the midline to innervate the contralateral ATL and SCL (Figures 4A and 4B). WPNs appear to have large output puncta in the ATL and SCL (Figures 4B and S3). We note that these neurons appear to be members of a large anatomical class that project along the same characteristic tract from the WED dorsally, crossing the midline at the level of the ATL and SCL (Figure S3E). By recording from other neurons in this class, we found some that were tuned (like the WPNs) for ipsilateral wind, though with a smaller dynamic range, while others responded only transiently to wind with no direction tuning (Figure S2). Thus, the two neurons labeled in the driver line *70B12-GAL4* are likely members of a larger mechanosensory projection to the dorsal brain.

**Figure 4.**
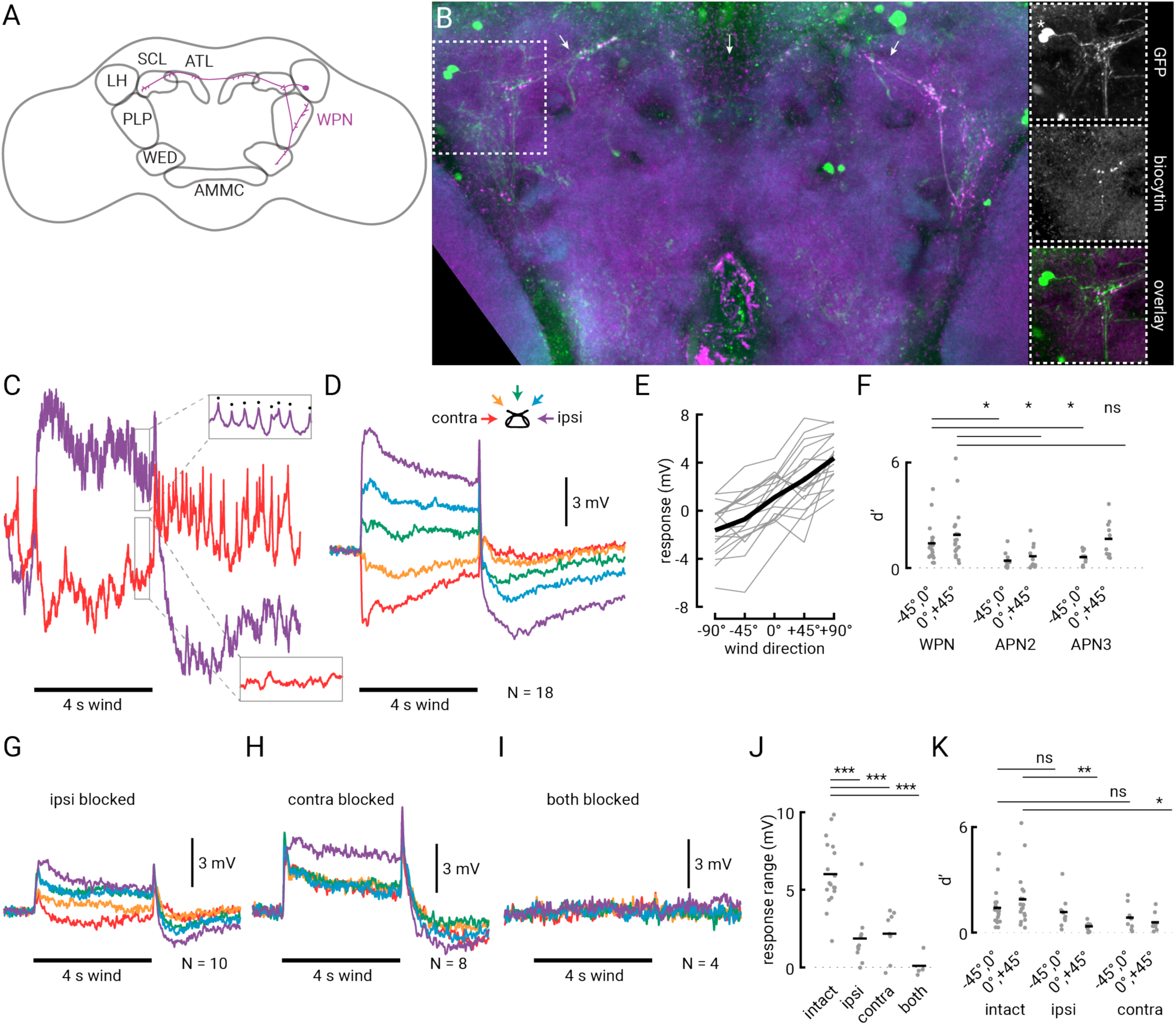
Wedge projection neurons combine input from the two antennae to linearly encode wind direction. (A) Schematic of a single wedge projection neuron (WPN) with processes in the WED, PLP, SCL and ATL. (B) Maximum z-projection showing two WPNs filled on the right hemisphere of the brain (magenta) overlaid with GFP (green) from the driver line used to target these cells (*70B12-GAL4*). Neuropil is shown in blue (left image only). Dotted white box indicates 60×60 μM region of interest expanded on the right. Arrows indicate axon tract that runs across the brain (see also Figure S2E). Top right: GFP signal only. Two cell bodies on the left side of the brain are indicated by the asterisk. Middle right: contralateral intracellular fill(s) in the expanded region. Note proximity of biocytin-filled puncta from ipsilateral (right hemisphere) cells and processes of contralateral cells labeled with GFP. Bottom right: overlay of GFP and biocytin in the expanded region. (C) Raw intracellular responses from one WPN to a 4 s ipsilateral (purple) and contralateral (red) wind stimulus. The neuron produces small-amplitude spikes at rest and during ipsilateral wind (upper box, spikes indicated by the black dots) and is inhibited by contralateral wind (lower box). (D) Average membrane potential of N=18 WPNs (from 17 flies) during wind stimuli from the five directions (schematic depicted above the traces). Same scale as (C). (E) Average steady state response (difference between 1 s at the end of the wind stimulus and the 1 s baseline immediately before wind stimulus is presented). Individual WPN averages in grey; average across 18 cells indicated by the thicker black line. (F) Discriminability index (d’) between responses to wind delivered from frontal directions (−45° vs. 0° and 0° vs. +45°) for WPN, APN2 and APN3. Single fly average d’ values are plotted as grey dots, and the horizontal line indicates average across flies. Asterisks indicate if the WPN’s d’ values are different from the corresponding values in APN2 or APN3 (two-sample t-test, P = 0.012, 0.026, 0.040, and 0.68). (G) Average WPN response from N=10 flies with mechanosensory input from the ipsilateral antenna blocked. (H) Average wind response from N=8 WPNs in flies with contralateral antenna input blocked. (I) Average responses from N=4 WPNs in flies with input from both antennae blocked. (J) Response range (difference between steady state ipsilateral and contralateral WPN wind response) for flies with free, ipsilaterally-blocked, contralaterally-blocked, or dual-blocked antennae. Single fly averages are shown as dots and the horizontal bar indicates the average across flies. Mean responses from flies with the ipsilateral, contralateral, or both antenna blocked are significantly different than flies with free antennae (two-sample t-test, P = 0.00022, 0.00012, and 0.000029, respectively). (K) Discriminability index (d’) between frontal directions (−45° vs. 0° and 0° vs. +45°) for WPNs with free antennae, or ipsilateral or contralateral input blocked. Single fly averages are shown as dots and the horizontal bar indicates the average across flies. Asterisks indicate if the WPN’s d’ values are different from the corresponding d’ values in flies with blocked antenna (two-sample t-test, P = 0.55, 0.0046, 0.21, and 0.030). All sample sizes for time traces are noted to the right of the black stimulus bars.

To characterize the wind responses of the WPNs, we presented wind to the fly from five directions and measured their responses using whole cell electrophysiology. WPNs were excited by wind delivered from the ipsilateral side and inhibited by wind from the contralateral side (Figures 4C-E). Distinct from the APNs, they exhibited an even gradient of tuning to intermediate wind angles (Figures 4D and 4E). WPNs produced small-amplitude spikes at a low baseline firing rate (8.4 Hz) and responded to tonic wind stimuli with both graded changes in membrane potential and changes in firing rate (Figures 4C and D, Figure S4). WPNs also exhibited a greater ability to differentiate between wind angles in front of the fly (0° vs 45° ipsilateral or contralateral) than APNs for most angles, as measured by the d-prime statistic (Figure 4F). Single neurons had somewhat differing levels of inhibition; however, the dynamic range of individual cells was similar (Figure 4E).

The striking linearity of the WPN’s wind response across azimuthal angles (Figure 4E), and the greater discriminability in front of the fly (Figure 4F), were both reminiscent of the difference between displacements of the two antennae (Figure 2F). Thus, we wondered whether WPNs integrate information across the two antennae. To address this question, we blocked input from either the ipsilateral (Figure 4G) or contralateral antenna (Figure 4H), and recorded wind responses in the WPN. Both manipulations greatly diminished directional wind responses in the WPN but did not abolish them, indicating that WPNs receive directional wind information from both antennae. Stabilizing or removing either antenna had statistically indistinguishable effects (Figure S5) and led to a significant reduction in dynamic range (Figures 4G, 4H, and 4J), as well as significant reduction in discriminability of some neighboring frontal angles (Figure 4K). Further, when we blocked input from both antennae, the WPN no longer responded to wind (Figure 4I), indicating that the WPNs receive all their input from the antennae. These results indicate that WPNs perform integration across the two antennae to generate a more linear representation of wind direction in azimuth and to increase the discriminability of frontal wind angles.

### Wedge projection neurons are not synaptically connected

We next aimed to identify where information from the two antennae is exchanged. By visualizing the processes of single WPNs filled during recordings (see Methods), we noticed that the outputs of these neurons overlap closely with the processes of WPNs on the contralateral side of the brain (Figure 4B). Therefore, we hypothesized that WPNs on opposite sides of the brain might synapse onto each other to exchange wind direction information. To test this hypothesis, we performed dual whole cell patch clamp recordings from pairs of WPNs on opposite sides of the brain (Figure 5). When we injected current into one neuron, we observed an increase in the rate of spiking in that neuron, but saw no response in the contralateral WPN, and vice versa (Figure 5A). We observed a complete absence of response in three pairs of contralateral WPNs in different flies (Figure 5B), supporting the notion that the WPNs are not synaptically connected.

**Figure 5.**
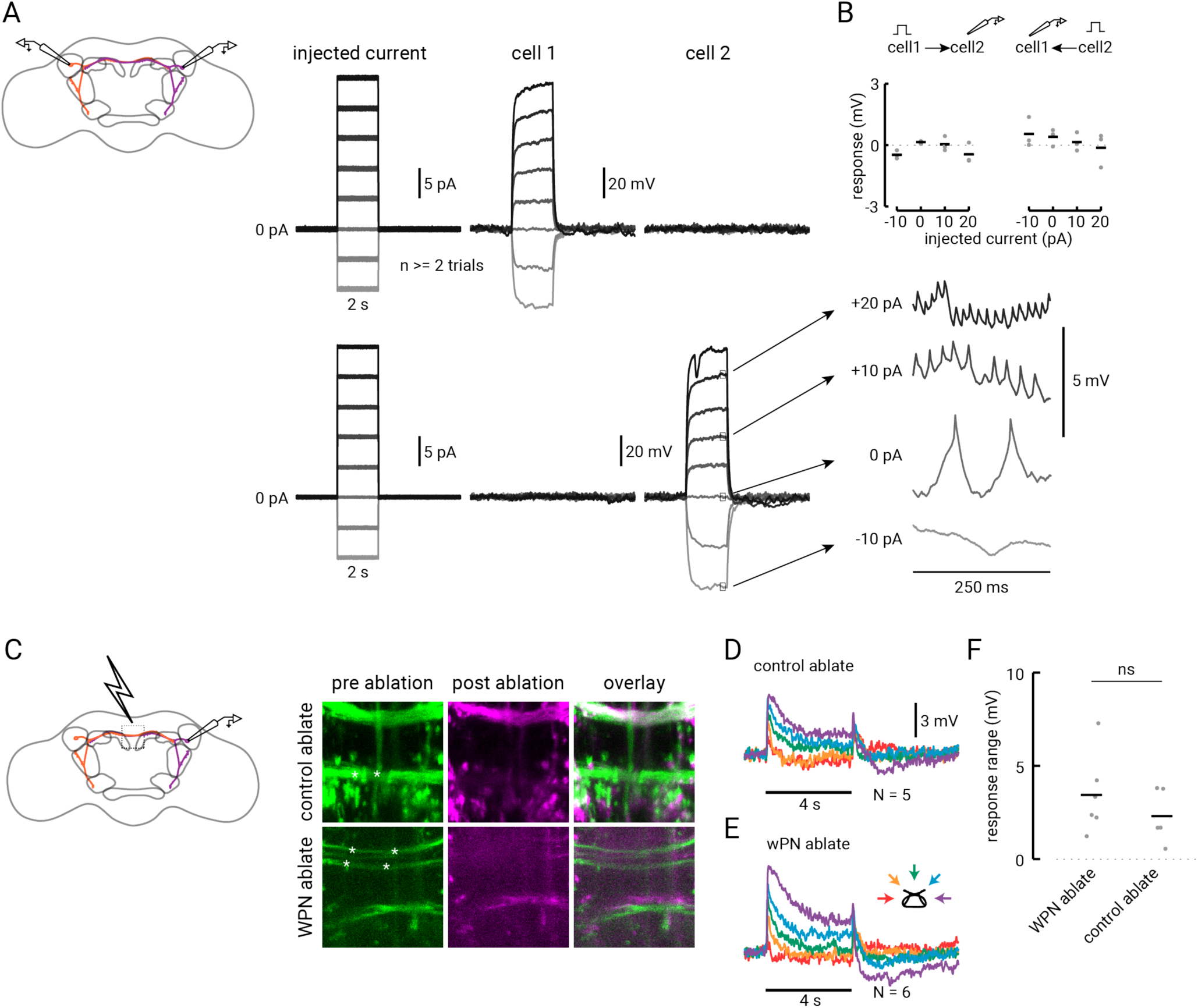
The contralateral arm of the WPN does not contribute to its wind tuning. (A) Dual whole cell recording from two WPNs on opposite sides of the brain (depicted in purple and orange in the schematic to the left). On the right, top traces show current injected into cell 1 (cell body located in the right hemisphere) and resulting response in the contralateral cell 2 (left hemisphere). Below traces show current injected into cell 2 and resulting response in cell 1. Expanded traces on the right show typical changes in spike rate due to current steps in that neuron. (B) Average response of contralateral partner to current pulses for three pairs of simultaneously recorded WPNs. None of the average responses across cells are significantly different than zero (one-sample t-test, from left to right, P = 0.056, 0.077, 0.89, 0.26, 0.33, 0.24, 0.65, 0.83). (C) Ablation of WPN or control axons using a 2-photon laser light pulse. Left: central region where axons were targeted for lesioning. Right: Example z-projections of a 41.18 μM square central region of the brain where either a control set of axons (top row; 30 μM deep) or the WPN axons (bottom row; 20 μM deep) are lesioned. Left, pre-ablation GFP signal from *70B12-GAL4*, with focal point of laser indicated by the white asterisks. Center: post-ablation stack of the GFP signal in the same region (magenta). Right: overlay of pre- and post-ablation images. (D) Average wind response of N=5 WPNs for flies in which we lesioned control axons. (E) Average wind response for N=6 WPNs with severed axons. (F) Mean steady state difference between ipsilateral and contralateral wind response in flies with either WPN or control axons lesioned. Average for the two conditions are not significantly different (two-sample t-test, P = 0.34).

One concern we had regarding this experiment was that the WPNs have very fine and extended processes, so current injection at the soma might not significantly affect activity at distant synapses. So in a complementary experiment, we used a high-energy 2-photon light pulse to lesion the contralateral projections of the WPNs and measured their resulting wind tuning (Figure 5C). As a control, in separate experiments, we lesioned a set of axons labeled in the same line that cross the midline more ventrally and do not appear to have overlapping processes with the WPNs. Figure 5 shows images from two examples in which we ablated either the control axons or the WPN axons in the center of the brain. We found that wind responses in WPN-lesioned and control-lesioned animals were similar (Figures 5D-F). Both groups showed reduced inhibition, but this reduction was not different between the two groups (Figure 5F; two-sample student’s t-test, P = 0.34), suggesting that it might arise from factors related to the lesioning protocol (see Methods). Together, the negative results from the dual patch and laser ablation experiments support the idea that information exchange between WPNs does not contribute to their wind tuning. Thus, we predict that WPNs receive information from the contralateral antenna from some other contralaterally-projecting neuron, perhaps at the level of the wedge.

### Multiple AMMC projection neurons contribute to WPN wind tuning

Because the WPN’s wind tuning does not appear to depend on its contralateral projections, we next asked whether integration across the antennae occurs at WPN inputs in the wedge. To address this question, we first generated a *70B12-lexA* driver line, and verified that the same two neurons are labeled by this line (Figure S3). Next, since other second-order mechanosensory neurons have been shown to receive large inputs through electrical synapses (Azevedo & Wilson, 2017), we asked whether wind responses in WPNs depend on chemical synaptic transmission. We blocked excitatory chemical transmission using the acetylcholine receptor antagonist methyllycaconitine (MLA, 1:1000) and found that this completely abolished the steady state wind response in WPNs (Figures 6A and 6B). This experiment indicates that the tonic WPN responses require excitatory chemical synaptic transmission.

**Figure 6.**
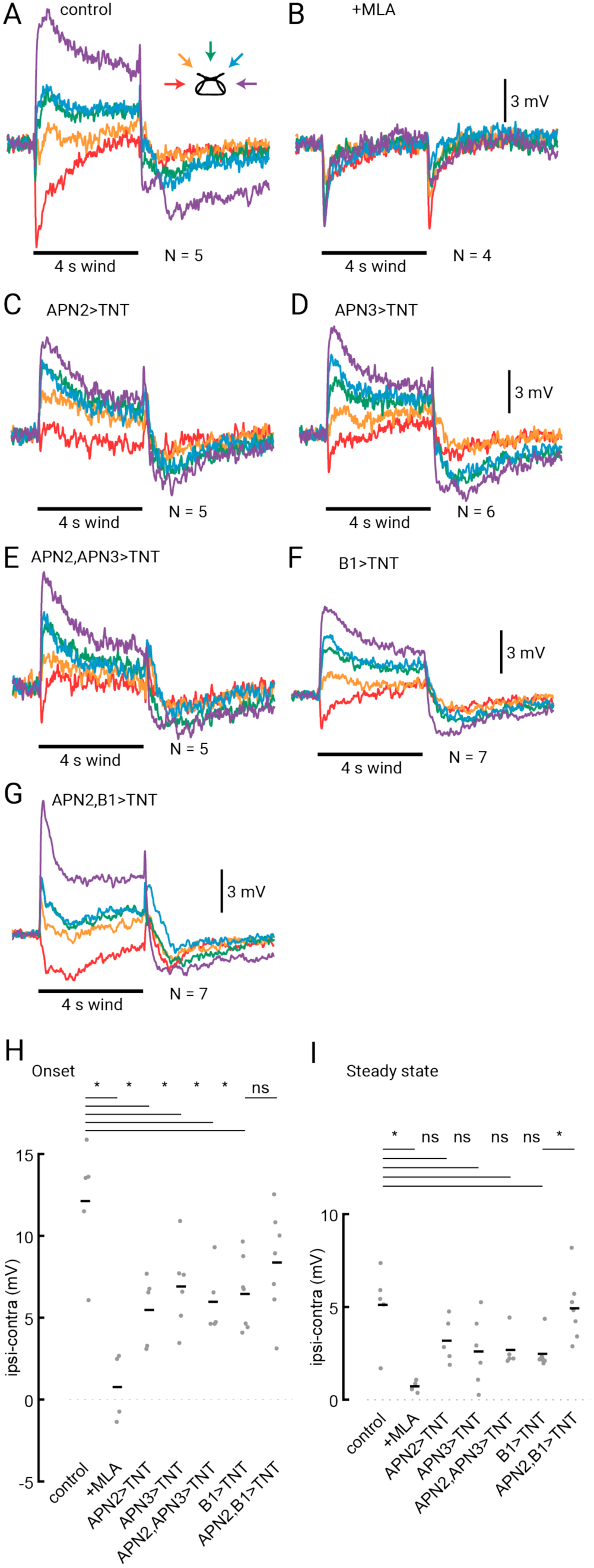
WPNs receive input from multiple AMMC projection neuron populations. (A) Average responses of WPNs to a 4 s wind stimulus. (B) Average wind response in WPNs after bath-application of methyllycaconitine (MLA; 1:1000), which blocks acetylcholine receptors (thus excitatory chemical synaptic transmission). (C) Average wind responses from WPNs in flies in which APN2 is silenced by expressing tetanus toxin (TNT). (D) Average WPN wind responses in flies with silenced APN3 neurons. (E) Average wind responses from WPN neurons in flies where both APN2 and APN3 are silenced. (F) Average WPN wind responses in flies with silenced B1 neurons. (G) Average WPN wind responses in flies in which both APN2 and B1 are silenced. (H) Mean difference between ipsilateral and contralateral WPN responses at wind onset (average over 800 ms near the start of the wind stimulus) in individual flies (gray dots) and across flies (horizontal black lines). Expression of TNT in any of the AMMC projection neurons (APN2, APN3, and B1) and in both APN2 and APN3 results in a significant change in mean response range (two-sample t-test; from left to right, P = 0.0010, 0.0085, 0.022, 0.012, and 0.0075). Silencing APN2 and B1 does not result in a significantly different WPN response range than when silencing B1 alone (P=0.22). (I) Average difference between ipsilateral and contralateral steady state wind response in WPNs for conditions in a-f. (from left to right, P = 0.0046, 0.11, 0.066, 0.047, 0.012, and 0.0057). Number of flies contributing to each average trace in (A)-(G) is listed to the right of the wind stimulus bar.

Next, we proceeded to inactivate putative presynaptic partners of the WPNs by driving expression of tetanus toxin, which blocks chemical synaptic transmission. To test the contribution of input from APN2 and APN3, we silenced each of these cell types (Figures 6C and 6D) and found that in each case the wind response in WPNs was significantly diminished. This effect was most pronounced in the onset response to wind (Figure 6H), however, we also observed a decrease in the tonic, or steady state, wind response in some cases (Figure 6I). Silencing APN2 and APN3 simultaneously also produced a decrease in wind tuning but did not abolish wind responses (Figures 6E, 6G and 6H). This result indicates that at least one other class of neuron must provide input to the WPNs.

What other neurons might provide input to WPNs? One compelling candidate are the B1 neurons. These AMMC projection receive input from the antenna ipsilateral to their cell bodies (Azevedo and Wilson, 2017), and project both ipsilaterally and contralaterally. They have been thought to be mostly auditory, based on their anatomical position downstream of the sound-sensitive A/B JONs (Kamikouchi et al., 2009; Vaughan et al., 2014), and the finding that silencing these neurons significantly impairs courtship song responses in females (Vaughan et al., 2014). However, a recent study (Azevedo & Wilson, 2017) showed that these neurons respond directionally to displacements of the antenna induced by a piezoelectric probe, and that a subset of these are tuned for low frequencies, suggesting a possible role in wind direction encoding. To test the hypothesis that B1 neurons contribute to WPN directional responses, we silenced B1 neurons, and recorded wind responses in WPNs. We found that this resulted in a diminished WPN wind response (Figures 6F-H).

In addition, we asked whether optogenetic activation of APNs and B1 neurons could drive responses in WPNs (Figure S6). We found that light alone could depolarize WPNs (Figure S6A). Optogenetic activation of APN2 with CsChrimson however produced a depolarization with a distinctly different time course: a transient depolarization, followed by a hyperpolarization, (Figure S6B-D) compared to genetic controls. APN3 activation produced no significant effect (Figure S6E-F). We then found that optogenetic activation of B1 neurons resulted in a transient hyperpolarization of WPN membrane potential (Figure S6G-H)— the opposite of what we observed with APN2 activation. Although the effects of activation were small, they are consistent with the idea that APN2 and B1 neurons may provide input, directly or indirectly, onto WPNs with opposite sign.

Curiously, silencing of B1 and APN2 neurons together resulted in a milder deficit than silencing either population alone (Figure 6G). This result argues that additional populations of neurons must provide directional input to WPNs. We therefore used trans-synaptic tracing to investigate what additional neural populations might play a role in wind processing.

### Trans-synaptic tracing suggests a larger circuit participating in wind processing

To broadly investigate circuits that might play a role in wind processing, we used *trans-Tango* (Talay et al., 2017) to identify neurons downstream of primary wind-sensitive mechanoreceptor neurons (JO-CE neurons labeled by *JO31-GAL4*) as well as the APN2, APN3, and B1 neurons labeled by each of our drivers. Consistent with previous reports, we found that JO-CE trans-Tango labeled both APN2 and APN3 as postsynaptic partners (Figure 7A, Video S1). In addition, this technique labeled at least two prominent commissural populations, including what we believe to be the B1 neurons and a second, more posterior tract with posterior cell bodies (Figure 7A) that may include AMMC-A1 and/or AMMC-Bb (Figure 7A; Matsuo *et al*., 2016b) and aLN (GCI; Vaughan et al., 2014). These findings support the idea that B1 neurons receive information from “wind” JONs and may participate in wind direction encoding as well as auditory processing. We also observed some labeling of WPN-like neurons, suggesting that WPNs may receive input directly from JONs or from another neuron labeled in this line with expression in the WED (Figure 7A, inset).

**Figure 7.**
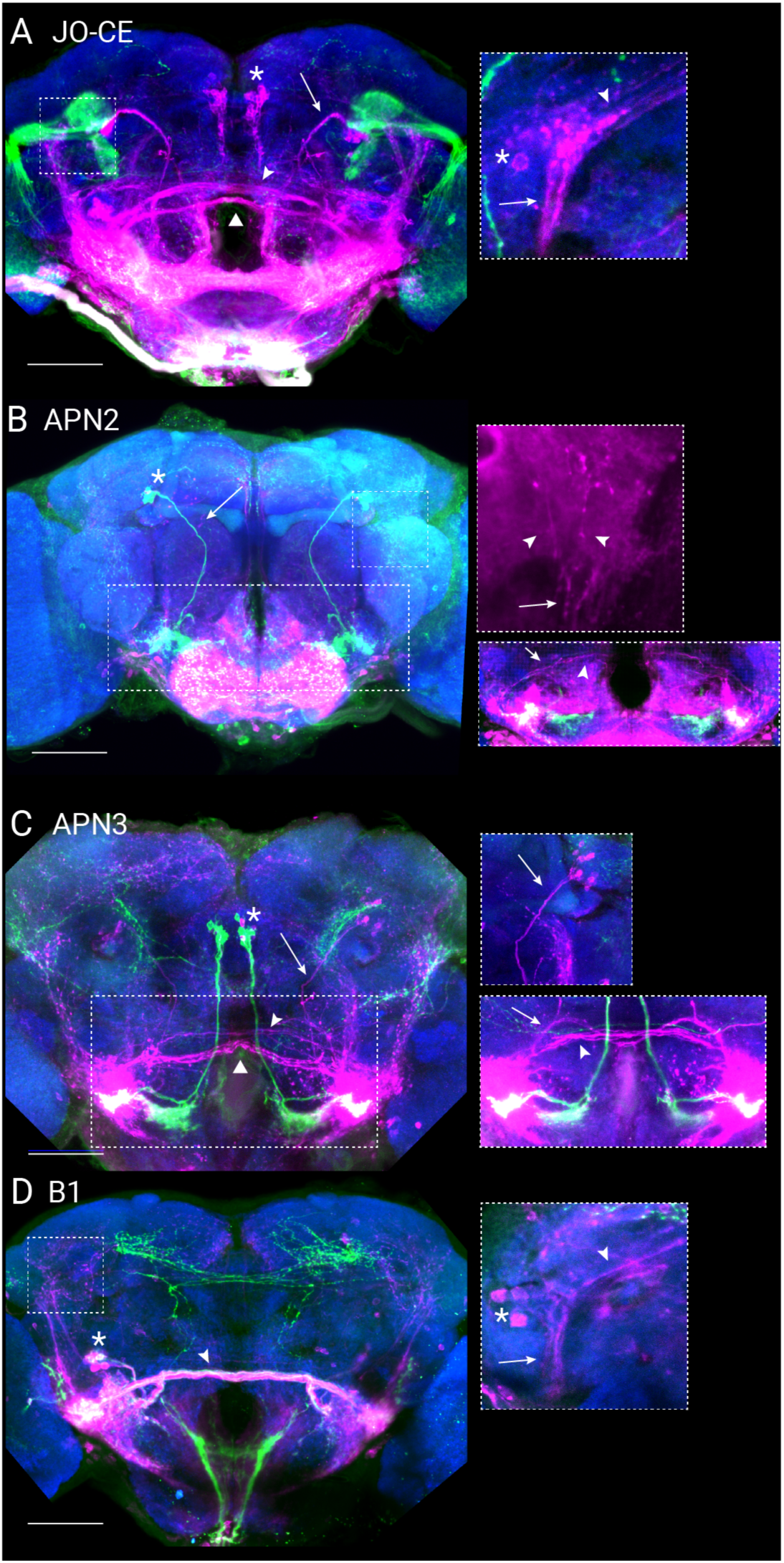
Trans-synaptic tracing labels multiple channels for wind encoding. (A) trans-Tango driven by primary wind mechanoreceptors (*JO31-GAL4*, green) labels putative postsynaptic neurons (magenta). Neuropil is shown in blue. Multiple second-order wind neurons classes are labeled, including APN2 (arrow indicates axon tract), APN3 (asterisk indicates location of cell body cluster), B1 (arrowhead points to axon tract passing the midline), and an additional commissural axon tract (triangle). Scale bar is 50 μM. Inset is a maximum projection of putative postsynaptic WPN-like processes (50×50 μM in size and 10 μM deep). Asterisk indicates cell body leading to WPN-like processes, arrow indicates axons leading town towards WED, and arrowhead indicates axons leading towards characteristic contralateral branch. (B) *trans*-Tango using APN2 (*24C06-GAL4*) as the presynaptic driver. Same colors and scale as (A). Upper right: Partial maximum projection (expanded 50×50 μM region, 14 μM deep, postsynaptic label shown alone for clarity) reveals WPN-like axon tract that leads dorsally out of the wedge (arrow) and branches into two distinct tracts in the PLP (arrowheads) Lower right: 10 μM deep maximum projection reveals B1 neurons (inset is 70×200 μM). Arrow indicates branching point of axons leading to the AMMC and WED, and the arrowhead indicates its contralaterally-projecting central axon. (C) trans-Tango using APN3 (*70G01-GAL4*) labels APN2 (arrow indicates axon tract; also see inset to the right), B1 (arrowhead), and an additional commissural axon tract (triangle). Asterisk shows location of cell bodies of APN3, some of which are self-labeled. Upper right inset: z-projection (100 μΜ square, 26 μΜ deep) showing post-synaptically-labeled APN2 cell bodies and axon (marked by arrow) leading down towards the WED. Same colors and scale as (A). Lower right inset: narrower z-projection (10 μΜ deep) highlights the B1 neurons (arrowhead) and an additional cell type leading to the WED (arrow; same scale as left image). (D) B1-driven *trans-*Tango. Same colors and scale as (A). Inset shows maximum projection of putative postsynaptic WPN-like processes (50×50 μΜ in size and 6 μΜ deep). Asterisk indicates cell body leading to WPN-like processes, arrow indicates axons leading town towards WED, and arrowhead indicates axons leading towards characteristic contralateral branch.

Next, we used trans-Tango to trace putative postsynaptic targets of APN2 (*24C06-GAL4;* Figure 7B). This labeled WPN-like processes, which can be identified most clearly where the dorsal axon exiting the wedge sends a major branch off into the posterior lateral protocerebrum (Figure 7B, inset). APN2>*trans*-Tango also labeled the same two commissures as in the JO-CE *trans*-Tango, however we believe this labeling arises from a cluster of small cell bodies near the B1 cell bodies that are also labeled in *24C06-GAL4* (Video S2). In contrast, APN3>*trans*-Tango did not appear to label WPNs. Some diffuse labeling was present along the region that runs between the WED and the posterior lateral protocerebrum (PLP), but these processes were much more anterior than the WPN axon tract between these structures (Figure 7C, Video S3). These results support our conclusions from silencing and optogenetic activation (Figure 6 and Figure S6), namely that APN3 neurons may connect only indirectly to WPNs. APN3>trans-Tango did label many putative second-order mechanosensory neurons, including APN2, B1, AMMC-Db1, and at least one other commissure (Figure 7, Video S3). This diversity of putative downstream partners suggests that APN3 broadly interconnects mechanosensory circuitry and may account for their paradoxical contralateral wind tuning. Finally, we analyzed the *trans*-Tango signal driven by B1 neurons. We observed no clear labeling of APN2 or APN3, but other putative third-order neurons appear, including distinct WPN-like processes at the posterior face of the brain and the corresponding cell body cluster (Figure 7D), consistent with the hypothesis that B1 neurons provide input to WPNs.

Together, our physiological and anatomical results suggest that wind direction is computed in the dendrites of WPNs at the level of the WED. At a minimum, this computation likely involves ipsilateral antenna deflection information, carried by APN2 neurons, and contralateral antenna deflection information, carried by B1 neurons (Figures 8A and 8B). However, our double silencing results (APN2 and B1 neurons) argue that WPNs must receive additional sources of directional information. Likely candidates are the neurons the project contralaterally in the posterior brain that we observed in our JO-CE transTango experiment (AMMC-A1, AMMC-Bb, and aLN). WPNs may also receive direct input from CE-JONs (Figure 7A). Finally, it is unclear whether these inputs synapse directly onto our two WPNs, or whether they synapse onto WPN-like neurons that pass information to neurons with similar projections. Future studies using connectomic approaches to reconstruct the full complement of inputs onto WPNs will be necessary to fully understand the circuits that allow WPNs to compute wind direction from displacements of the two antennae.

**Figure 8.**
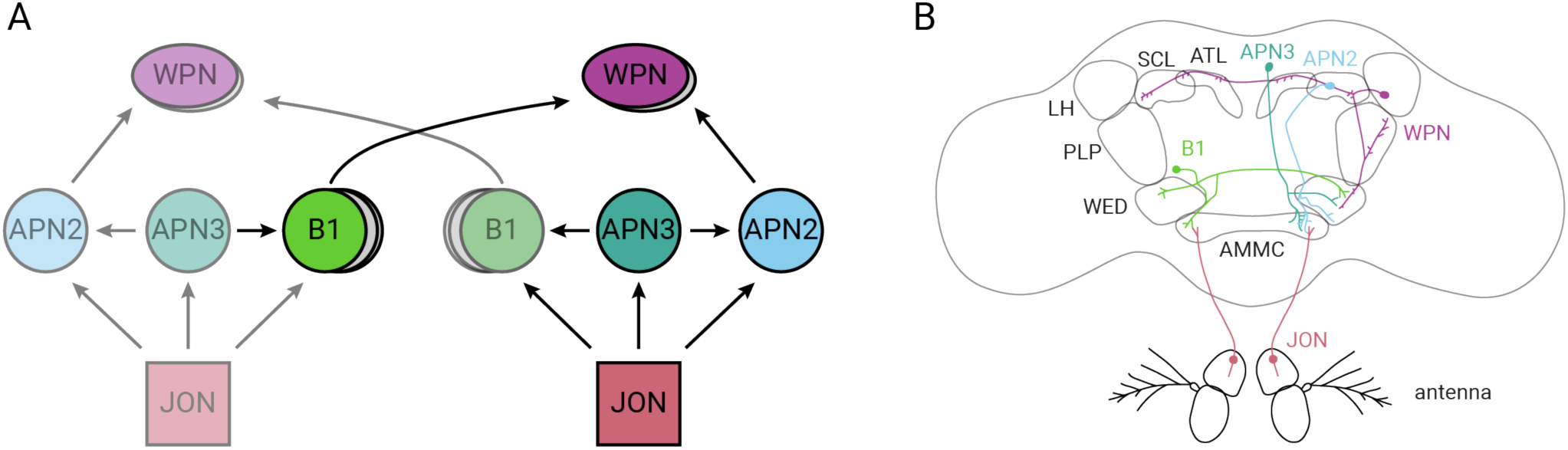
Summary diagram of the wind processing circuit. (A) Simple circuit diagram of wind neurons in this study, using previously established nomenclature for brain regions (Ito et al., 2014). Arrows indicate hypothesized direction of information flow. (B) Schematic depicting anatomical projection patterns for single neurons from the classes included in (A) (same colors). Neuropil regions are outlined in gray.

## DISCUSSION

### Wind direction as a mechanosensory computation

Animals use a wide array of sensory cues to navigate through the world. Visual cues are often used for navigation, because they can provide fast, reliable information about an animal’s location in space. However, mechanosensory cues can also provide spatial information. Animals that hunt in the dark, such as owls and bats, rely on highly specialized mechanosensory systems to compute the location of prey from sound stimuli. Studies of these systems have revealed many basic principles of neural computation (Konishi, 2003).

Wind is one such mechanosensory cue used by animals to navigate through space. When wind direction is stable, it can provide an orientation cue that permits dispersal over long distances (Bell & Kramer, 1979), or it can serve as a compass to help locate a feeding site or nest (Müller & Wehner, 2007). Odor molecules are typically dispersed by wind; consequently, many animals have evolved strategies of using wind direction cues, in combination with olfactory cues, to locate food or mates (David, Kennedy, & Ludlow, 1983; Flugge, 1934; Kennedy & Marsh, 2004; Lacey, Ray, & Cardé, 2014; Wolf & Wehner, 2000). In flying animals, orientation to wind is thought to require measurements of optic flow (Kennedy 1941; Kennedy and Marsh 1974; Murlis et al. 2000), because wind displaces the entire animal as well as its mechanoreceptors. In contrast, walking animals can derive wind direction directly from mechanosensation (Álvarez-Salvado et al., 2018; Bell & Kramer, 1979; Wolf & Wehner, 2000). Although this strategy, known as “odor-gated anemotaxis,” has been studied most extensively in invertebrates, there is some evidence that suggests that vertebrates use this strategy as well. For example, rats can use their vibrissae to determine wind direction (Yu, Graff, Bresee, Man, & Hartmann, 2016), and wolves will orient upwind after scenting their prey (Mech, 1966).

Like sound localization, computing wind direction also requires the nervous system to combine information from multiple sensors. A well-studied wind-sensing system is the cercal system, a set of paired appendages on the abdomen of crickets and other orthopteran insects. The cercus is covered with an array of bristles, each of which has a preferred axis of deflection. The cercal bristles create an array of direction-tuned axonal fields that are sampled by interneurons in the abdominal ganglion. These interneurons generate a new array of cosine-tuned wind sensors, each aligned with a few cardinal axes (Boyan and Ball, 1990; Jacobs, Miller, Aldworth 2008).

In contrast, the neural basis of wind-sensing by the antennae has been less studied, despite their critical role during odor-gated wind orientation in many walking insects (Álvarez-Salvado et al., 2018; Wolf & Wehner, 2000). In this study, we showed that flies require signals from both antennae to perform full upwind orientation during walking olfactory navigation, although a single antenna is sufficient to perform partial upwind orientation. We saw no effect of single antenna stabilization on downwind preference in the absence of odor. This may be because our measurement of downwind preference reflects the animals’ integrated behavior over time, or because flies are able to use active movements of the body, or of the antenna, to compensate for the loss of one sensor. Flies can actively position their antennae using muscles that innervate the 1^st^ antennal segment (Mamiya et al. 2011) and could use such movements to gain additional information about wind direction. We measured a slight increase in angular velocity during the odor stimulus, suggesting that flies might use body scanning to compensate for the loss of one antenna, and observed qualitative examples of looping. However, future studies will be necessary to fully understand how flies combine active movement with antennal mechanosensation during wind orientation.

To understand the anatomical basis of this wind sensitivity, we performed a detailed quantification of how the two antennae are displaced by wind from different directions and found that the geometry of the antennae underlies a fly’s ability to encode wind. More specifically, we found that a single antenna provides an ambiguous signal about wind direction because it is maximally deflected by wind roughly perpendicular to the arista, and displacement falls off symmetrically with deviations from this angle. Previous work (Morley et al., 2012) showed that the mechanical sensitivity of the antennae are greatest in the frontal field at 45° relative to the midline; indeed, we observe the largest antennal deflections when wind is delivered at −45° and 0° contralateral to each antenna. The central neurons we characterized in this study encode either the displacement of one antenna (with the associated ambiguity), or a signal closely resembling the difference between the two antennal displacements. The geometry of the antennae varies greatly across insect evolution (Krishnan & Sane, 2015); for instance different fly species from the same superfamily have distinct antennal morphologies (Sinclair & Cumming, 2006). How these different geometries might impact wind perception and sensitivity is an open question.

### Wind circuits in the fly brain

Previous studies hypothesized that antennal displacement signals must be integrated to form a central representation of wind direction (Patella & Wilson, 2018; Yorozu et al., 2009). Wind direction is first represented by tonically responding JONs (Kamikouchi et al., 2009; Yorozu et al., 2009), and functional imaging from JON axon terminals has shown that the E region of the AMMC responds to deflections of the ipsilateral antenna towards the head (“push”), while the C region responds to deflections away from the head (“pull”; Kamikouchi *et al*., 2009; Yorozu *et al*., 2009). Unlike in the mosquito (Andres et al. 2016), *Drosophila* JONs do not receive synaptic input (Kamikouchi et al. 2010). Thus integration is unlikely to occur at the level of JONs. Recently, pan-neuronal imaging identified a region of the WED that responds selectively to ipsilateral pull and contralateral push, the displacement pattern produced by ipsilateral wind (Figure 2, Patella & Wilson, 2018). However, this study did not identify specific neurons that perform these computations, or identify higher-order areas of the brain that might be involved in wind processing.

Here we have identified a novel class of neurons, which we call WPNs, that integrate displacement information from the two antennae. These neurons are excited by ipsilateral wind and inhibited by contralateral wind, resulting in a rather linear encoding of wind direction in azimuth. In addition, responses to nearby frontal wind angles are well-differentiated from one another. Both these coding features closely resemble the difference in displacement signals we measured across the two antennae during wind stimulation and are distinct from the tuning exhibited by either single antenna. To directly test whether WPNs perform integration, we recorded their responses in flies with stabilized antennae, and found that stabilization of either antenna reduced wind tuning and discriminability of frontal angles, consistent with integration.

To localize the site of integration we performed paired recordings from WPNs, 2-photon lesions of WPN processes, and recordings, activations and silencing experiments of putative upstream partners of WPNs. Taken together these experiments support a model in which information from the two antennae is integrated in the WED, by combining ipsilateral information carried by APN2 neurons with contralateral information carried by B1 neurons. However, our experiments also suggest that the circuit underlying the computation of wind direction is more complex than this simple model. First, we were unable to completely abolish directional wind responses by silencing any pair of inputs. This suggests that at least three and likely more populations carry wind direction information directly or indirectly to WPNs. At least two prominent contralateral tracts were labeled by trans-synaptic tracing downstream of wind-sensitive JONs. Some CE JONs are known to project contralaterally (Kamikouchi et al., 2009), and these might contribute to integrative responses in WPNs as well. Second, the effects of activating putative upstream partners (APN2 and B1 neurons) were generally weak and biphasic. This could indicate that connections between AMMC projection neurons and WPNs are indirect, going either through WED local neurons, or through other WPN-like neurons. Finally, we observed several paradoxical responses in the wind circuit. For example, APN3 neurons responded differently to stimuli that displaced the ipsilateral antenna by the same amount (−90° and +45°). In addition, we found that silencing both B1 and APN2 neurons resulted in milder deficits than silencing either population alone. Both of these findings suggest a role for recurrent connections that might amplify or modulate responses to antennal displacement. A comprehensive understanding of computations in the wind direction circuit will likely require EM reconstruction, which has very recently been accomplished for the well-studied olfactory projection neurons of the fruit fly (Frechter et al., 2018), as well as genetic tools for silencing larger numbers of GAL4 lines simultaneously.

### Potential downstream targets and functions of wind neurons

The higher-order wind neurons described in this study, WPNs, project to multiple targets in the brain, suggesting a broad dispersal of wind information throughout the dorsal brain. These regions include the posterior lateral protocerebrum (PLP), superior clamp (SCL), and antler (ATL). Although morphologically similar regions of the brain have been described in other insects (Immonen, Dacke, Heinze, & el Jundi, 2017; von Hadeln, Althaus, Häger, & Homberg, 2018), to our knowledge, little is known about the function of these regions of the brain. Neurons in the central complex of other insects have been shown to respond to mechanosensory stimuli (U. Homberg, 1994; Uwe Homberg, 1985; Ritzmann et al., 2008), but it is not fully clear how information from the APNs or WPNs might be routed to the central complex. Each brain region innervated by the wind neurons in our study (AMMC, WED, PLP, SCL, ATL; see Figure 8) is also the site of input for descending neurons (Hsu & Bhandawat, 2016; Namiki, Dickinson, Wong, Korff, & Card, 2017), raising the possibility that different features of wind information are relayed to motor systems multiple times. The identification of WPNs as mechanosensory neurons provides a starting point for investigating the function of these areas of the insect brain.

How might the wind direction information encoded by the WPNs be used during behavior? Because WPNs are activated by ipsilateral wind, they could produce upwind orientation if they activate neurons that drive ipsilateral turns. Conversely, they could produce downwind orientation by activating neurons that drive contralateral turns. The contralateral projections of the WPNs, which we found did not contribute to their wind-tuning, might carry information to such motor systems. WPNs may also participate in other forms of orientation such as detecting body orientation relative to gravity (Kamikouchi et al., 2009) or measuring the direction from which a song is produced during courtship behavior (Morley et al., 2012).

In our study, we have shown how information about wind direction is processed starting from the periphery and eventually integrated by higher order neurons that transmit information across the central brain. We also provide evidence that points to a site of mechanosensory information exchange from the two sides of the brain. Together, these results reveal novel pathways and encoding strategies for wind information and highlight the importance and prevalence of wind information in the *Drosophila* brain.

## ACKNOWLEDGEMENTS

The authors would like to thank Nicholas Stavropoulos for helping us generate the *70B12-lexA* flies, Matthieu Louis for *UAS-TNTe* flies, Mustafa Talay and Gilad Barnea for the *trans-Tango* flies, and Allison Baker-Chang and Rachel Wilson for the *R70G01-GAL4, 10xUAS-CD8:GFP* fly stock. We also thank Gerry Rubin for sharing Flylight expression data and Heather Dionne and Gerry Rubin for the R70B12 plasmid. We thank Dimitry Rinberg for use of his 3D printer and for critical feedback on this work. Greg Suh also provided critical feedback on the manuscript. Henry Haeberle help set up the Thorlabs software for 2-photo lesion experiments. This work was funded by NIH R00DC012065, NIH R01MH109690, NSF IOS-1555933, the Klingenstein Foundation, the McKnight Foundation, the Sloan Foundation (K.I.N.), and the Leon Levy Foundation (M.P.S.).

## AUTHOR CONTRIBUTIONS

M.P.S. and K.I.N. designed antenna tracking and electrophysiology experiments. M.P.S. performed and analyzed electrophysiology experiments. M.P.S., S.S. and M.D. analyzed videos and antenna tracking data. A.M.M.M., K.I.N. and M.P.S. designed behavior experiments; A.M.M.M. and M.P.S. conducted and analyzed the behavior experiments. A.M.M.M., K.I.N., and M.P.S. acquired and analyzed confocal images. M.P.S., K.I.N., and D.S. designed the laser ablation experiments. M.P.S. performed and analyzed laser ablation experiments. M.P.S. and K.I.N. wrote the manuscript with critical input from all co-authors.

## KEY RESOURCES TABLE

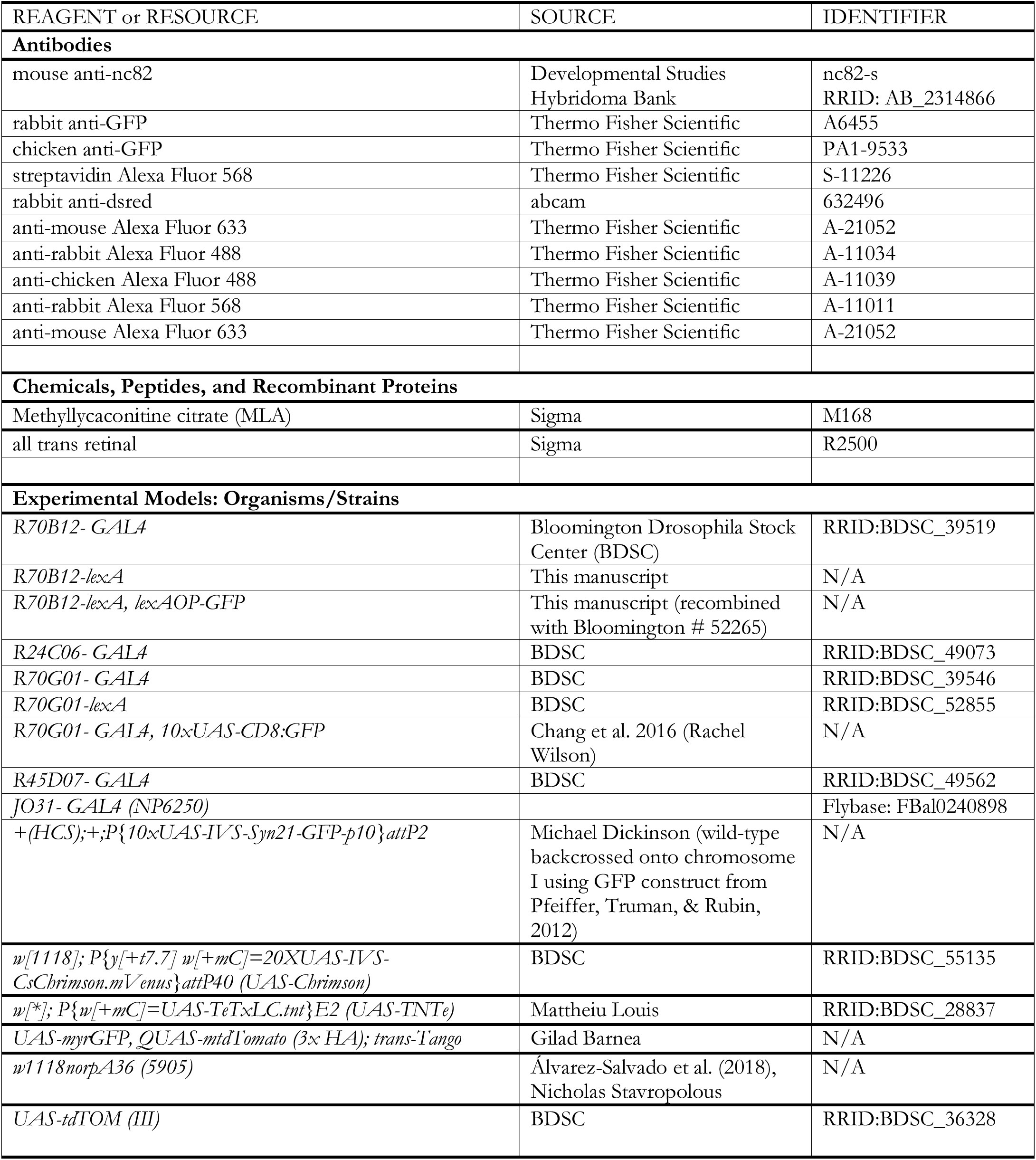
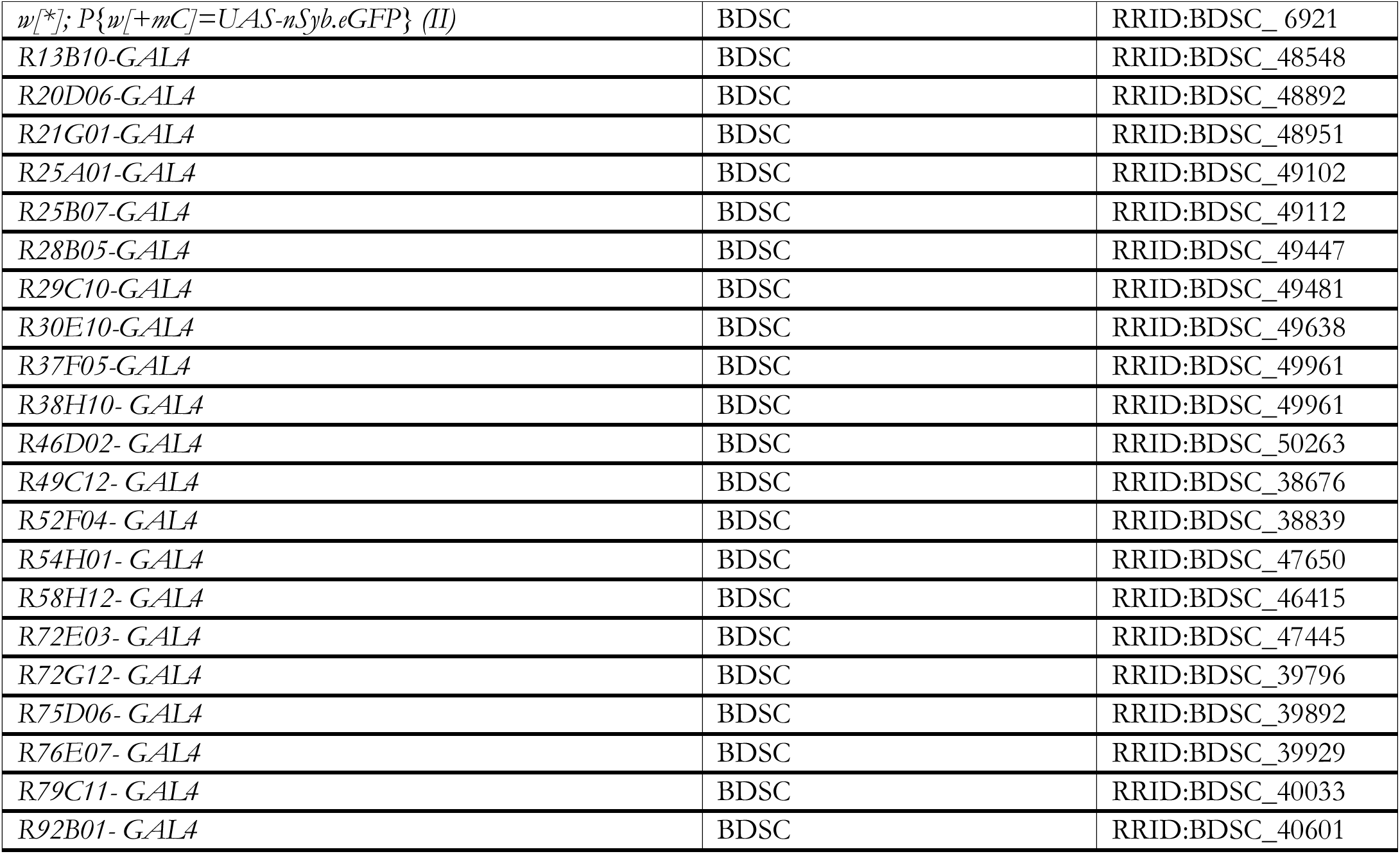

Contact for reagent and resource sharing: Katherine Nagel

## METHODS

### Fly strains

For behavioral experiments, we used flies bearing *norpA^36^* backcrossed into an isogenic *w^1118^* strain (Bloomington #5905) that exhibits robust walking and olfactory searching behavior (Álvarez-Salvado et al., 2018). To target cell bodies for electrophysiological recordings, we expressed cytoplasmic GFP (typically *10XUAS-IVS-Syn21-GFP-p10* but also *10xUAS-CD8:GFP*) under control of the following driver lines: *R24C06-GAL4* (APN2), R70G01-*GAL4* (APN3), and *R70B12-GAL4* (WPN). We made flies carrying *R70B12-lexA* using standard techniques (Pfeiffer et al., 2008; Pfeiffer et al., 2010). This was recombined with *lexAOP-GFP* (from Bloomington #52265) to visualize WPNs while using the GAL4-UAS system to silence putative upstream neurons. For Chrimson activation experiments, we used flies with the following genotypes: *70B12-lexAop, lexAop-GFP / UAS-CsChrnmson; R24C06-GAL4/UAS-10xGFP* and *R70B12-lexAop, lexAop-GFP/UAS-CsChrimson; R70G01-GAL4/UAS-10xGFP*. For inactivation experiments, we used the three following genotypes: *R70B12-LexAop, LexAop-GFP/UAS-TNTe; R24C06-* GAL4and *R70B12-lexAop, lexAop-GFP/UAS-TNTe; R70G01-GAL4*, and *R70B12-lexAop, lexAop-GFP/UAS-TNTe*; R45D07-*GAL4*. Control flies were *R70B12-lexAop, lexAop-GFP/+*. For trans-synaptic tracing using *trans-*Tango, we crossed *UAS-myrGFP, QUAS-mtdTomato (3x HA); trans-Tango* flies to *R24C06-GAL4* (APN2), *R70G01-GAL4* (APN3), *R45D06-GAL4* (B1) or *JO31-GAL4* (JO-CE neurons). In the text, we refer to all GAL4-driver lines from the Janelia Flylight collection without the ‘R’ prefix for simplicity. All the genotypes we in this study are listed in the following table.

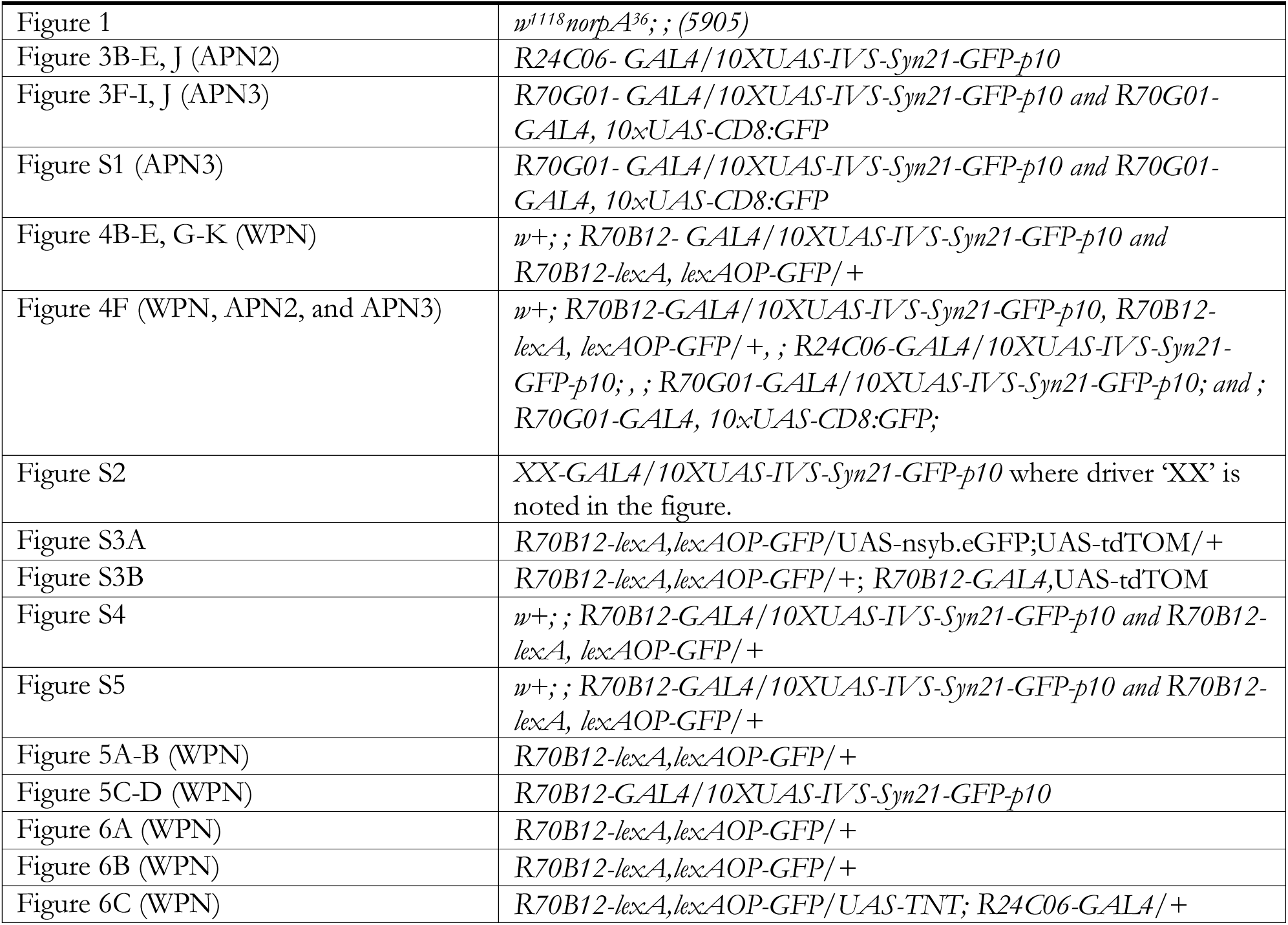
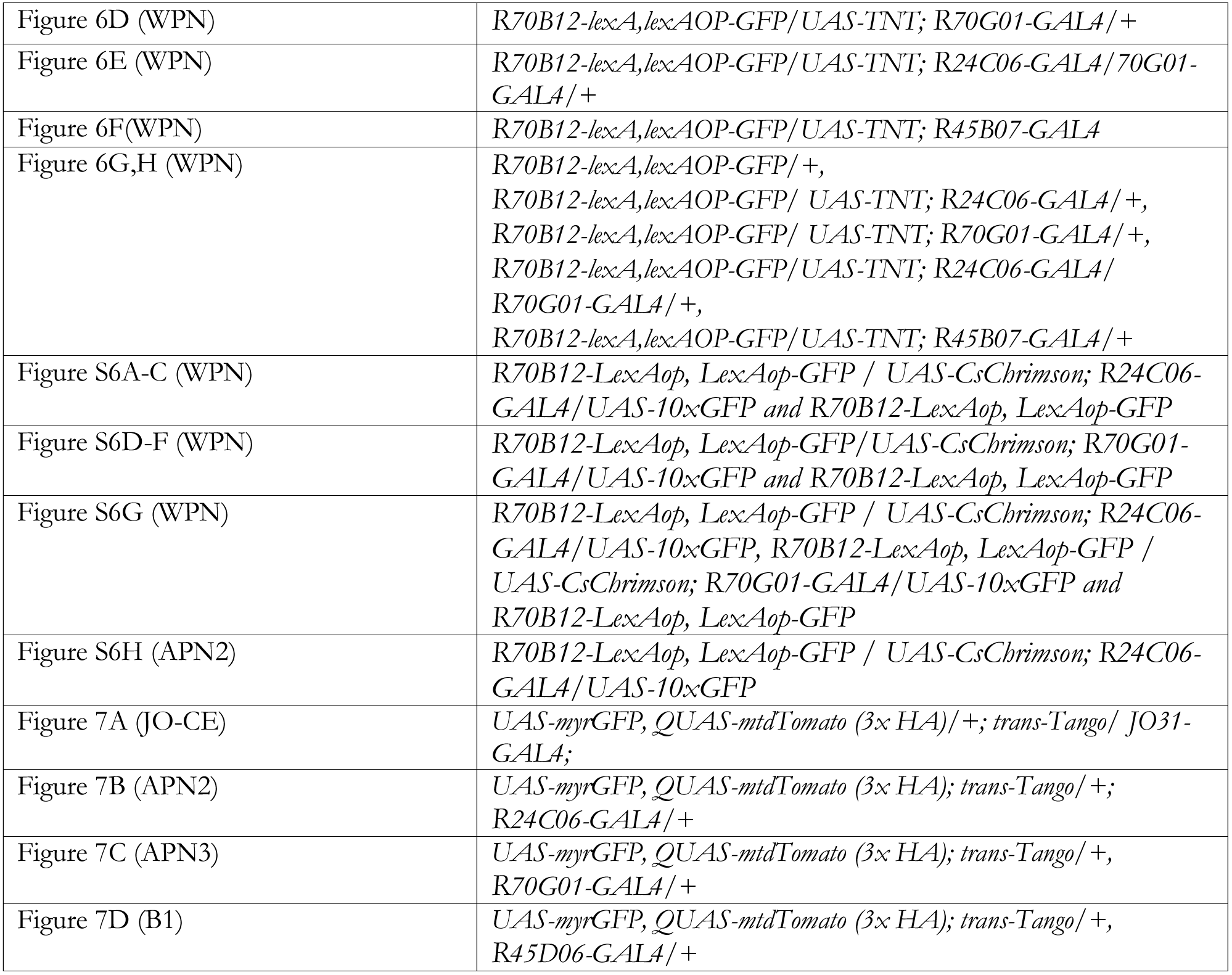

### Behavior

For behavioral experiments, we used the same miniature wind tunnel apparatus described in Álvarez-Salvado *et al*. (2018). As previously described, flies were constrained to walk within a shallow arena. A constant airflow travelled through the arena at 11.9 cm/s as measured by a calibrated hot wire anemometer. A 10 s pulse of odor (1% apple cider vinegar, Pastorelli) was introduced by a set of solenoid valves located immediately below the arena. Advection of the odor through the arena was measured by photo-ionization detector (miniPID, Aurora Scientific, Aurora Canada). Flies randomly experienced one of 3 conditions on a given trial: wind only, a 10 s odor pulse in constant wind (30 s wind, 10 s odor, 30 s wind) or no wind. We collected mated females 3–7 days before the experiment and housed them in custom time shift boxes on light cycles of 12 h. We deprived flies of food for 20–24 h before experiments in an empty polystyrene vial with a small amount of shredded damp Kimwipe on the bottom to humidify the air. Experiments lasted approximately two hours were started at either ZT1 or ZT3 (1 or 3 hours after lights on).

To block mechanosensory input from the Johnston’s organ in one or both antennae, we gently anesthetized a fly over ice, and removed the fly’s arista with a sharp pair of forceps by pinching it off precisely near its base. We then applied a small amount of UV glue (Aquaseal UV Field Repair Adhesive) to the antennal joint between the 2^nd^ and 3^rd^ segments near the ablated arista and cured for 10 s. We used a small drop of glue so that the majority of sensilla on the third segment were unobscured. Flies were allowed to recover for 20–24 h before the experiment, during which time they were deprived of food, but not water (similar to all other behavioral experiments). In a pilot set of experiments, we observed that the glue was no longer present on the antennae for some flies (we hypothesize that some were able to groom it off during the long starvation period). Thus, after each experiment, we gently removed each fly from the behavioral arena and inspected its antenna(e) to verify that the manipulation persisted throughout the starvation period and experiment. For control flies, we performed a “sham” manipulation in which we cold-anesthetized and exposed the flies to UV light for the same duration as experimental flies.

We analyzed behavioral data using similar protocols to those described in Álvarez-Salvado *et al*. (2018). Briefly, we tracked the position and orientation of flies in real time using custom Labview software (National Instruments) and further analyzed the data in Matlab (Mathworks). We discarded any trials with tracking errors, and low-pass filtered coordinates and orientations at 2.5Hz using a 2-pole Butterworth filter. Trials in which flies moved less than 25 mm, and flies that had fewer than 5 trials that met our criteria were discarded. In addition, we excluded trials in which flies spend more than 25% of the period used for analysis within 3 mm of the edge of the arena. Because odor reaches flies at different times depending on their position in the arena, we aligned trajectories for upwind velocity to the actual time the odor reached the fly based on their position and the delay as recorded by the PID (miniPID, Aurora Systems). When measuring angular velocity, we aligned trials to the end of the odor presentation. All flies that were excluded from the odor condition were subsequently excluded from the wind preference assay. One additional fly was excluded from the arena position analysis (Figure 1G) in both glue and single glue conditions compared to Figure 1B because of lack of movement during wind-only trials. Behavioral parameters (upwind velocity and absolute angular velocity) were computed only using segments in which flies moved at greater than 1 mm/s.

Upwind velocity and angular speed were computed for the 5 s following odor onset and the 2 s following the offset of the odor, respectively, on a fly-by-fly basis and averaged. We averaged behavior from 10 s after the trial began to 5 s before odor onset for a total of 15s to compute the baseline level of activity.

### Wind stimulus apparatus

We custom-designed the wind stimulus delivery apparatus (Autodesk Inventor Professional) to precisely align a mounted fly at the center of 5 wind streams of equal velocity. The apparatus was 3D printed (Makerbot Replicator 2) and consisted of 3 parts: the wind manifold, the physiology holder, and the base. The wind manifold had five symmetric wind channels with a diameter of 0.1352 in (a bit wider than a fly). The physiology holder was created based on previous designs (Maimon, Straw, & Dickinson, 2010; Weir et al., 2016) (see also http://ptweir.github.io/flyHolder/), and we designed the base to hold the manifold and the physiology holder at a constant angle and position. The manifold was attached to a base by a mounting screw which also allowed us to fix the apparatus to a post. The physiology holder attached to the base in a fixed position using magnets. When the fly was mounted, the openings of the channels were 0.55 in away from the fly’s head.

To generate the wind stimuli, we passed house air over a charcoal filter and a water reservoir (to humidify the air). Wind was controlled by a set of 5 solenoid valves (Lee Company LHDA1233115H) that directed air flow through one of the five channels, or to a vacuum. Wind and vacuum flow rates were set using flow meters (Cole-Parmer PMR1–010608). Valves were controlled from Matlab via a digitizer (National Instruments BNC-2090A) that interfaced with a microcontroller board (Arduino Mega 2560). The Arduino decoded control signals and controlled a set of relays to direct the wind flow through one of the five channels, or to the vacuum. We placed flyback diodes across the leads of the solenoid valves to eliminate a large voltage spike that we otherwise picked up with the recording electrode.

We measured wind velocity at the location of the fly using an anemometer (Dantec Dynamics MiniCTA 54T42) with a probe (Dantec Dynamics 55P16) placed in the position normally occupied by the fly. Windspeed at the position of a mounted fly was 59.57 (+/− 1.4) cm/s. By mounting the probe in parallel with the wind channels, such that the filament was perpendicular to this flow, we ensured that it was circularly symmetric with respect to the five channels. We mounted the probe 6 times in this way and averaged the results to obtain a realistic measurement of stimulus variability (e.g. resulting from small deviations in the fly’s size and/or position in the physiology holder). To calibrate the probe and calculate absolute wind velocity, we created a pseudo-laminar wind flow with a C-mount tube filled with coffee straws and a mass flow meter placed 20 mm from the anemometer probe. A mass flow meter (Aalborg Mass Flow Controller GFC17 0–2L) was then used to relate air velocity (computed by dividing the volume flow rate by the cross-sectional area of the tube) to the voltage readout of the anemometer. The linear range of this function (20 to 70 cm/s at the anemometer) was used to determine absolute wind velocity.

### Antenna tracking

To monitor movements of the antennae in response to wind, we acquired videos of the fly from below using an Allied Vision Guppy Pro F-031B camera with an Infinistix 68mm/2.00x lens with 90-deg mirror running at 60 Hz. We triggered the camera and acquired videos using Matlab. For all flies, we first defined the midline axis of the head using points at the top and bottom of each eye (at the edge of the retina) consistently across animals. To quantify antenna displacements over time (Figure 2D), we then used simple feature extraction techniques to detect the angle of the arista, relative to the midline of the head. Briefly, to detect the line of the main branch of the arista in each image, we performed the Hough transform of the image, found the peaks of the resulting Hough transform matrix, and detected line segments corresponding to these peaks. Errors were detected by excluding angles that were far out of the range of realistic antennal deflections (outside of the 30 deg range in which the antenna was displaced in these experiments). In the single fly tracking example shown in Figure 2D, we observed three instances in which there were large active deflections of the antenna; we omitted the trial block in which these occurred, to quantify the passive deflections of the antenna only. We averaged over 7 trials (7 videos) per direction for this example fly to obtain the average antenna deflection over time.

To quantify displacement as a function of wind direction (Figure 2E and 2F), we found that hand-tracking the antenna was the most accurate method for quantification. For these data, two people (M.D. and S.S.) were blinded to the wind direction in each trial. They hand-digitized the position of the arista in individual video frames using two points to define the line of the main branch. For each trial/video, we computed the average baseline arista position across 5 frames just before the wind stimulus turned on, and the steady state deflection across 5 points at the end of the wind stimulus. We did this for every fly, averaged across trials within a fly (over a minimum of 1 and maximum of 10 trials per direction, typically 5), then averaged across flies.

### Electrophysiology

We performed whole cell patch clamp recordings in tethered female flies using previously described methods and solutions (Maimon et al., 2010; Wilson & Laurent, 2005). Briefly, we cold-anesthetized flies and glued them to a plastic holder using UV-curing glue (Kemxert Corp York KOA 300). The holders were made by 3D printing a plastic base (MakerBot Replicator 2), and attaching metal cutouts (Etchit) to the holder using UV-curing clue using guidelines outlined in (Weir et al., 2016). We removed the front two legs at the base of the femur, and the rest of the legs at the base of the coxa, to prevent interference with the wind stimulus. We then applied UV glue to the proboscis, securing it to the head and to the front two coxa, to prevent excessive brain movement. To access the brain, under saline, we removed the cuticle at the back of the head using a small hypodermic needle (30G), and then using fine forceps, we gently removed only the trachea on the surface of the brain over the region where our cell bodies resided. For most flies, we also removed muscle 1 which would otherwise obscure cell bodies of interest. We broke through the sheath of the brain and cleaned the area only around the cell bodies of interest by using positive pressure to apply a small amount of collagenase (5% in extracellular saline; Worthington Biochemical Corporation Collagenase Type 4) with a fine-tip electrode (approximately 5–10 μM diameter at the tip). Extracellular saline, composed of 103 mM NaCl, 3 mM KCl, 5 mM TES, 8 mM trehalose dihydrate, 10 mM glucose, 26 mM NaHCO_3_, 1 mM NaH_2_PO_4_H_2_0, 1.5 mM CaCl_2_H_2_O, and 4 mM MgCl_2_6H_2_O, was continuously perfused over the brain during recordings.

For whole cell patch clamp recordings, we pulled 7–13 MΩ glass pipettes using a Sutter Instruments P-1000 puller (smaller resistance electrodes were used with the smaller WPN and APN3 cell bodies, and larger with the larger APN2 cell bodies). Our intracellular solutions contained 140 mM KOH, 140 mM aspartic acid, 10 mM HEPES, 1 mM EGTA, 1 mM KCl, 4 mM MgATP, 0.5 mM Na_3_GFP, and 13 mM biocytin hydrazide (for visualization of neural processes). The average membrane potential of the wind neurons were −22.3 mV, −22.0 mV, and −26.4 mV for WPNs, APN2, and APN3, respectively. These values are not adjusted for an estimated -13 mV junction potential measured in a previous preparation with identical solutions, preparations, and electrodes (Suver, Huda, Iwasaki, Safarik, & Dickinson, 2016).

We used the following three patch clamp amplifiers in our electrophysiology experiments: A-M Systems Model 2400 (for part of the wind neuron screen), Multiclamp 700B (Molecular Devices; for dual-patch and B1 recordings), and an Axopatch 200B (Molecular Devices) paired with additional preamplification by a Brownlee Precision Model 410 preamplifier for the remaining (majority) of experiments. We analyzed data and controlled stimuli with custom Matlab software. Control signals were relayed to stimulus devices via a digitizer (National Instruments BNC-2090A). We acquired all data at 10 kHz and then downsampled to 1kHz for analysis. Before downsampling, we detected spikes by smoothing (over a window of 20 samples), differentiating, and thresholding the membrane potential. False-positives were eliminated by excluding any spikes that occurred within 4 ms of each other, and the accuracy of this spike detection was confirmed by eye for each cell type. We filtered out spikes from the membrane potential averages for neurons that produce small spikes (APN3 and WPN) by lowpass filtering at 20 Hz. All average APN and WPN recordings are presented as averages across flies in which we obtained a stable recording for a minimum of 2 of each of the 5 wind directions, and a maximum of 15 trials/direction (7 trials on average). For all neural classes, we targeted a mixture of left and right-hemisphere cell bodies for recordings, and responses from neurons in the left hemisphere were flipped and averaged with those on the right. We targeted all APN2 neurons for recordings using *24C06-GAL4*, and APN3 neurons using *70G01-GAL4*. However, we targeted eight WPNs using cytoplasmic GFP under the control of *70B12-GAL4*, and ten using cytoplasmic GFP under control of *70B12-LexA*; we observed no difference between these groups.

### Statistics

For both the behavioral and electrophysiology data, we used the Jarque-Bera test determine if the data sets were normally distributed. The majority of data groups from the electrophysiology recordings were normal, thus we computed statistics for these experiments using standard parametric tests (two-sample student’s t-test) and corrected for multiple comparisons with the Bonferroni method. For the behavioral experiments, the majority of data sets were not normally distributed, so we used non-parametric statistics to compare these results. We performed a Wilcoxon signed rank test to compare the mean upwind velocity and absolute angular speed values across flies with the mean baseline value (Figure 1C and 1E). In Figure 1D and 1F, we subtracted the baseline from the response for each individual fly and compared between conditions by performing a Mann-Whitney U test on the baseline subtracted parameters. The positions of the flies used to generate the probability distributions in 1G were computed using the last 30 s of trials to account for the fact that the preceding trial can influence the arena position of a fly at the beginning of the trial. We computed probability distributions for each fly in a given condition and averaged these to generate the plots shown. We performed a Mann-Whitney U test on the average arena positions across flies to determine if there was a significant difference in position across the conditions in Figure 1G. In Figure 1C-G, we corrected the threshold for significance with the Bonferroni method. For all of the behavioral and electrophysiology data, we present standard deviation around mean values, with one exception. For Figure 1B, we depict standard error around the mean time traces, because standard deviation was large enough that it obscured the underlying average trend. For this data, however, we plot standard deviation and single-fly averages in subsequent subfigures (Fig. 1D-F) to fully depict the variability present in the data set. We computed the d-prime (d’, also called the discriminability index in Figure S2D) statistic in Figures 4K and S2 for pairs of directions for each neuron using the following equation:

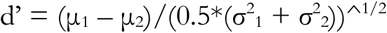

where μ_1_ and μ_2_ are the average steady state responses of the neuron for two adjacent wind directions, and σ^1^ and σ^2^ are their standard deviations, respectively, across trials. For Figure S2, we then averaged the d’ statistic across all neurons to compute the average d’ value.

### Pharmacology

To block cholinergic synaptic transmission, we added 1:1000 methyllycaconitine (1mM solution in dH20 stored at 4°C) to our extracellular saline.

### Optogenetic activation

For Chrimson activation experiments, we raised flies on typical agar food supplemented with 50 μL all-trans retinal (35mM stock dissolved in ethanol, stored at 4°C) mixed into 1 tablespoon of hydrated potato flakes. We targeted WPN neurons using 70B12-lexA and expressed UAS-CsChrimson under the control of either *24C06-GAL4* (for APN2 activation) or *70G01-GAL4* (for APN3 activation). We performed light control recordings in flies with the following genotype: *70B12-lexA/lexAop-10xGFP; 24C06-GAL4*. Responses elicited in APN neurons were relatively small (a little over 5 mV, see Figure S6), so we used the lowest light intensity that produced a maximum response from APN2 neurons (2.23 mW/cm^2^; Figure S6).

### 2-photon laser ablation

For lesioning, we mounted flies in the same way as for electrophysiology experiments (see *Electrophysiology* above). Immediately before lesioning, we removed the cuticle and trachea (again identically to electrophysiology experiments). The brain was submerged in fresh saline, but was not perfused, for approximately 20 min. Recording was delayed by about 50 minutes (on average) relative to recordings without laser ablation. We performed axotomies using an apparatus pioneered by Tsai et al. (2009) and optimized by Koyama et al. (2016). We used a high-power (8W) fixed wavelength (1040 nm) low-repetition rate (200 kHz) pulsed infrared laser (305 fs) with integrated pulse picker (Spirit One, Spectra-Physics). The high-power laser beam was concentrically aligned with a standard Ti-Sapphire two-photon imaging laser (Mai Tai DeepSea), steered using a ThorLabs microscope controlled with ThorImage 3.0, and focused on to the sample using a 40x objective (Olympus LUMPLFLN 40x/0.8 W). For each axon, we delivered 1–2 repeats of a 10-pulse long train at 10 kHz. Pulse power was 39 mW measured at the sample with a power meter (ThorLabs, PM 100D and S130C). We acquired pre-lesion z-stacks immediately before axotomy, and post-lesion z-stacks approximately 10 s after exposure to the high-power pulse train.

### Immunohistochemistry and anatomy

To visualize processes of recorded neurons, we gently removed the fly from the physiology holder and dissected the fly’s brain in PBS immediately after a recording. We then fixed the brain for 15 min in 4% paraformaldehyde solution (in PBS). Next, we washed the brain in PBS three times and stored at 4°C until antibody staining (within three days). We incubated brains in blocking solution (5% normal goat serum in PBST) for 20–60 min and incubated them for 24 hours at room temperature in a primary antibody solution containing 1:10 mouse anti-bruchpilot (nc82, Developmental Studies Hybridoma Bank) and 1:1000 rabbit anti-GFP (Thermo Fisher Scientific #A6455) in block. After three washes in PBS, we incubated the brains in a secondary antibody solution containing 1:250 anti-mouse Alexa Fluor 633 Thermo Fisher Scientific A-21052), 1:250 anti-rabbit Alexa Fluor 488 (Thermo Fisher Scientific A-11034), and 1:1000 streptavidin Alexa Fluor 568 (Thermo Fisher Scientific S-11226) in block, again at room temperature for 24 hours. We washed brains three times in PBS and mounted them in vectashield (Vector Labs H-1000). Slides were imaged immediately or stored at 4°C until imaging. We imaged brains at 20x magnification on a Zeiss LSM 800 confocal microscope with 20x objective (Zeiss W Plan-Apochromat 20x/1.0 DIC CG=0.17 M27 75mm). All brains were imaged at 1 μM depth resolution. The final images presented in our figures are maximum z-projections.

To define neuropil regions of interest in Figure 8 and Figure S2E, we traced these regions on small overlaid z-projections each at different depths corresponding to the structure. We further consulted standard brain maps to guide these boundaries relative to other structures in the brain (VirtualFlyBrain.org and Ito et al., 2014).

For *trans-Tango* images and for comparing expression of *70B12-GAL4* and *70B12-lexA* drivers, we used the above immunohistochemistry and imaging protocol, except that we used a primary antibody solution containing 1:50 chicken anti-GFP (Thermo Fisher Scientific PA1–9533), 1:500 rabbit anti-RFP (abcam ab62341) and secondary solution containing 1:250 anti-chicken Alexa Fluor 488 (Thermo Fisher Scientific A-11039) and 1:250 anti-rabbit Alexa Fluor 568 (Thermo Fisher Scientific A-11011), and anti-mouse Alexa Fluor 633 (Thermo Fisher Scientific A-21052) in block. We raised trans-Tango flies at 18°C and aged them until they were 10–20 days old.

**Figure S1 (related to Figure 3).**
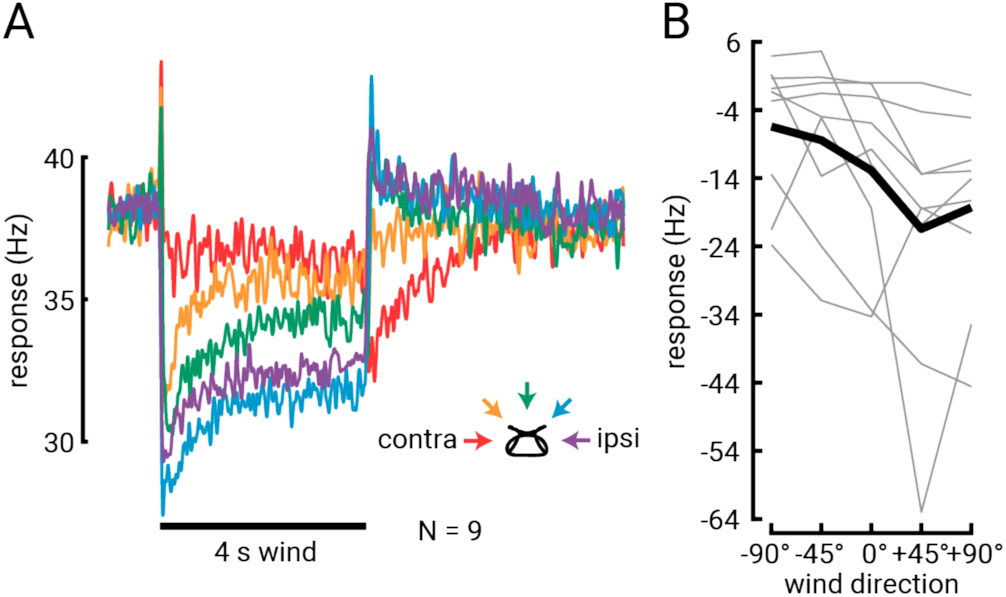
Spiking responses of APN3 neurons to wind. (A) Average spike rate of N=9 APN3 neurons in response to the 4 s wind stimulus from five directions. (B) Steady-state tuning of single APN3 neurons (thin lines) and across all 9 neurons (thick black line). Averages are derived from the last 1 s of wind stimulus region relative to the 1 s baseline before the wind stimulus is turned on.

**Figure S2 (related to Figure 4).**
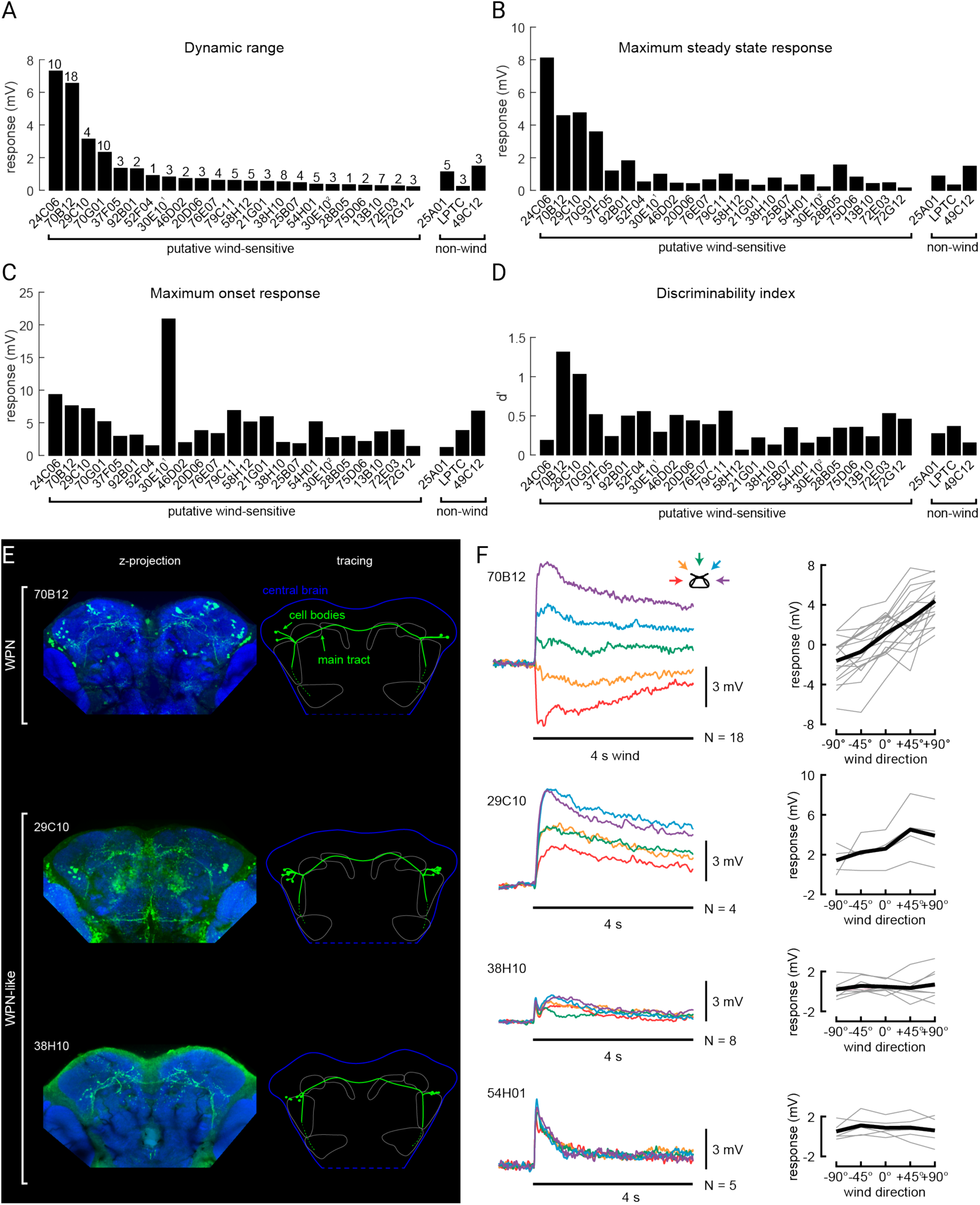
Wind direction neuron screen. (A)-(D) Response measures for each cell type targeted, including both candidate wind lines and non-wind controls (right). The responses are ordered based on descending response range, and the name of each GAL4 driver is plotted at its response value (except for lobula plate tangential neurons [LPTCs], which we targeted in more than one driver based on stereotypic size, position, and response to moving gratings). The same distinct group or pair of posterior cell bodies were targeted in each driver, except for *30E10-GAL4*, in which we targeted either one distinct large pair of neurons (30E10^1^), or a nearby set of smaller cell bodies (30E10^2^). All putative wind-sensitive drivers were chosen based on their processes in the WED, ATL, or PLP (nearest to the LH); three classes of neurons were analyzed for comparison: lateral horn intrinsic neurons (*25A01-GAL4*), lobula plate tangential cells (LPTCs), and a pair of mushroom body output neurons (*49C12-GAL4*). (A) Dynamic range (maximum-minimum responses) for average steady state wind responses. (B) Average maximum steady state response across the wind directions. (C) Average maximum wind onset response. (D) Frontal wind direction discriminability index (average across −45° vs. 0° and 0° vs. +45°) for target neurons in each driver. Averages shown in (A)-(D) are for the same N=X neurons for each neuron class, where X is listed above the average in (A). (E) Anatomy of WPN and “WPN-like” cell classes. The left column shows maximum z-projections across 10–20 μM sections of each brain to highlight the main axonal tract of the neuron. The right column shows a rough tracing of this main tract, and location of the cell bodies of the neurons targeted for recordings (arrows). An outline of the central brain is shown in blue (the bottom portion of each brain is cut off in this image, and indicated by the dashed blue line). The portion of the neurons that extend out of the PLP and into WED, as well as the major brain regions where these neurons project to, are not fully apparent in the maximum projection shown here; they have instead been traced from the same brain at different z-depths (dashed green and solid grey lines). (F) Left: Average wind response of WPNs and neurons with “WPN-like” anatomy labeled by three driver lines. 4 s wind stimulus indicated by black bar, same colors as main text. Right: Average steady state wind tuning for individual cells (grey lines) and across flies (black line) for each of the four driver lines from the left.

**Figure S3 (related to Figure 4).**
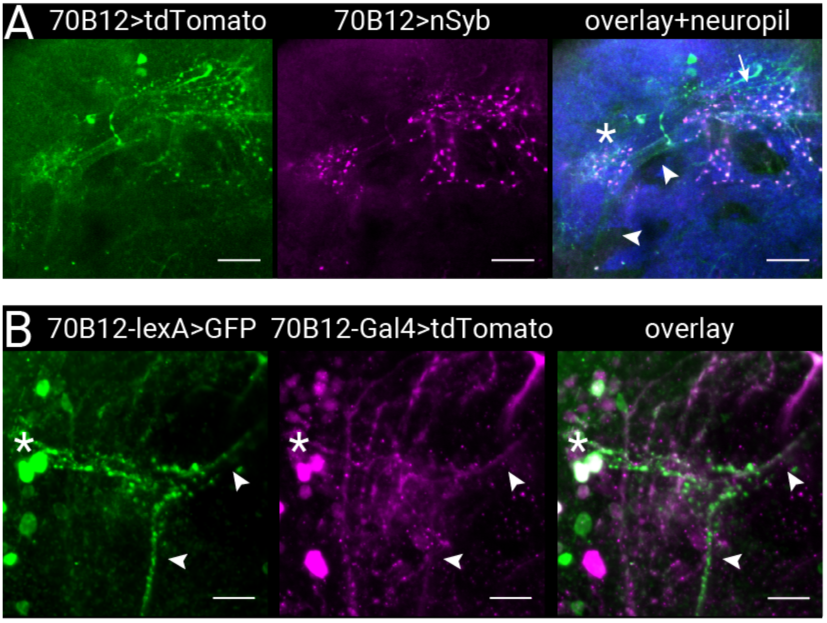
Polarity of neural processes in WPNs and validation of WPN-lexA driver. (A) WPNs have output processes in the SCL and ATL. Expression of tdTomato, driven by the genetic line we use to label WPNs (*70B12-GAL4*) is shown in green. Pre-synaptic staining driven by nSyb.eGFP is shown in magenta. Overlay, along with neuropil stain (blue) is shown on the right. Output puncta in the SCL are marked with an asterisk, and in the ATL by the arrow. WPN axons leading up to the SCL, and from the SCL towards the ATL are indicated by arrowheads. Scale bar is 10 μM. (B) The same two WPN cell bodies are labeled by *70B12-GAL4* and *70B12-lexA*. Confocal z-projection (16 μM deep) showing expression in WPN cell bodies by driven by *70B12-GAL4* (red) and *70B12-lexA* (green). Cell bodies in the left hemisphere are shown (marked by the asterisk) and appear white due to overlap. Distinct WPN axons leading towards the SCL are noted by the arrowheads. Scale bar is 10 μM.

**Figure S4 (related to Figure 4).**
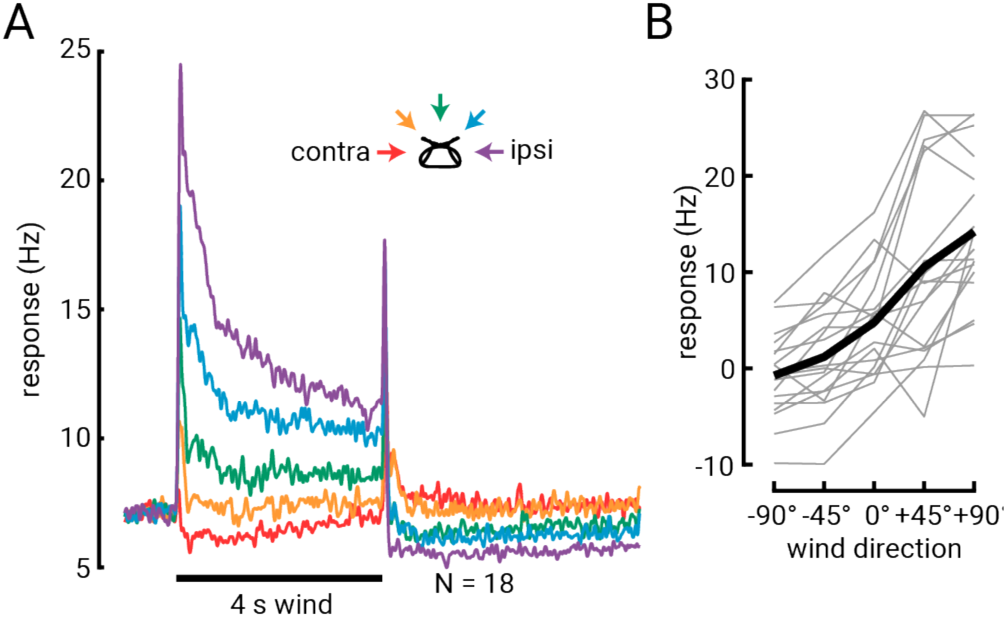
WPN spike rate responses to wind stimuli. (A) Average spike rate of WPNs in response to a 4 s wind pulse. Average across 18 flies (derived from same data as plotted in Figure 4d). (B) Steady state tuning curve (difference between 1 s at the end of the wind stimulus and the 1 s baseline immediately before wind stimulus is presented) for spike rate data plotted in (A).

**Figure S5 (related to Figure 4).**
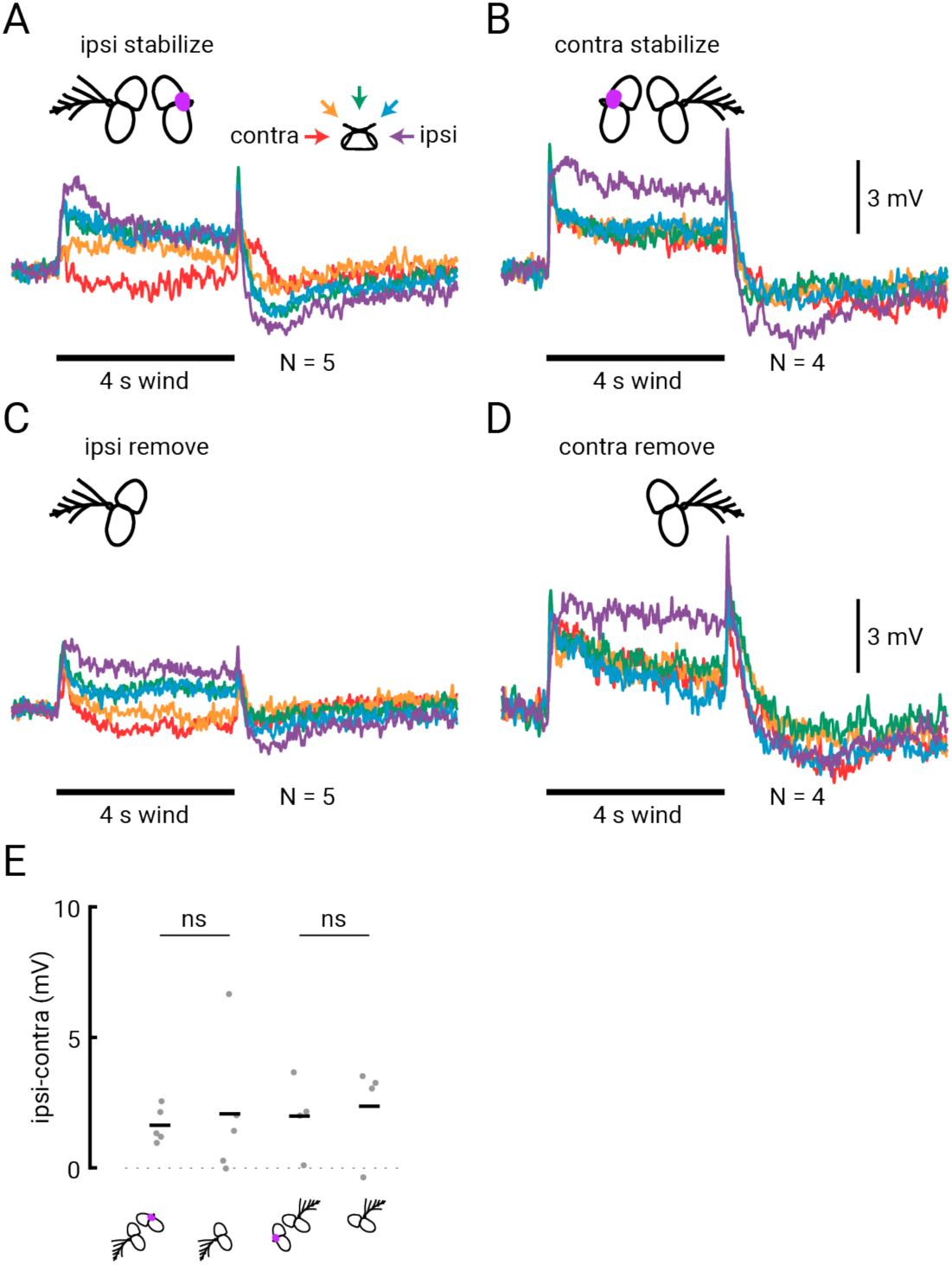
WPN wind responses are similar when one antenna is stabilized versus removed. (A) Average wind responses from WPNs in flies in which we cut the arista at its base and blocked movement of the third segment of the ipsilateral antenna with glue. (B) Average WPN responses from flies in which we cut the arista and stabilized the third segment of the contralateral antenna. (C) Average WPN responses from flies in which we removed the 2^nd^ and 3^rd^ segments of the ipsilateral antenna. (D) Average responses from WPNs in flies with contralateral antenna removed. (E) Difference between steady state ipsilateral and contralateral wind responses in flies with stabilized versus removed antenna. The means between ipsilaterally- or contralaterally-stabilized versus removed antenna are not significantly different (two-sample t-test, P = 0.74 and 0.76, respectively).

**Figure S6 (related to Figure 6).**
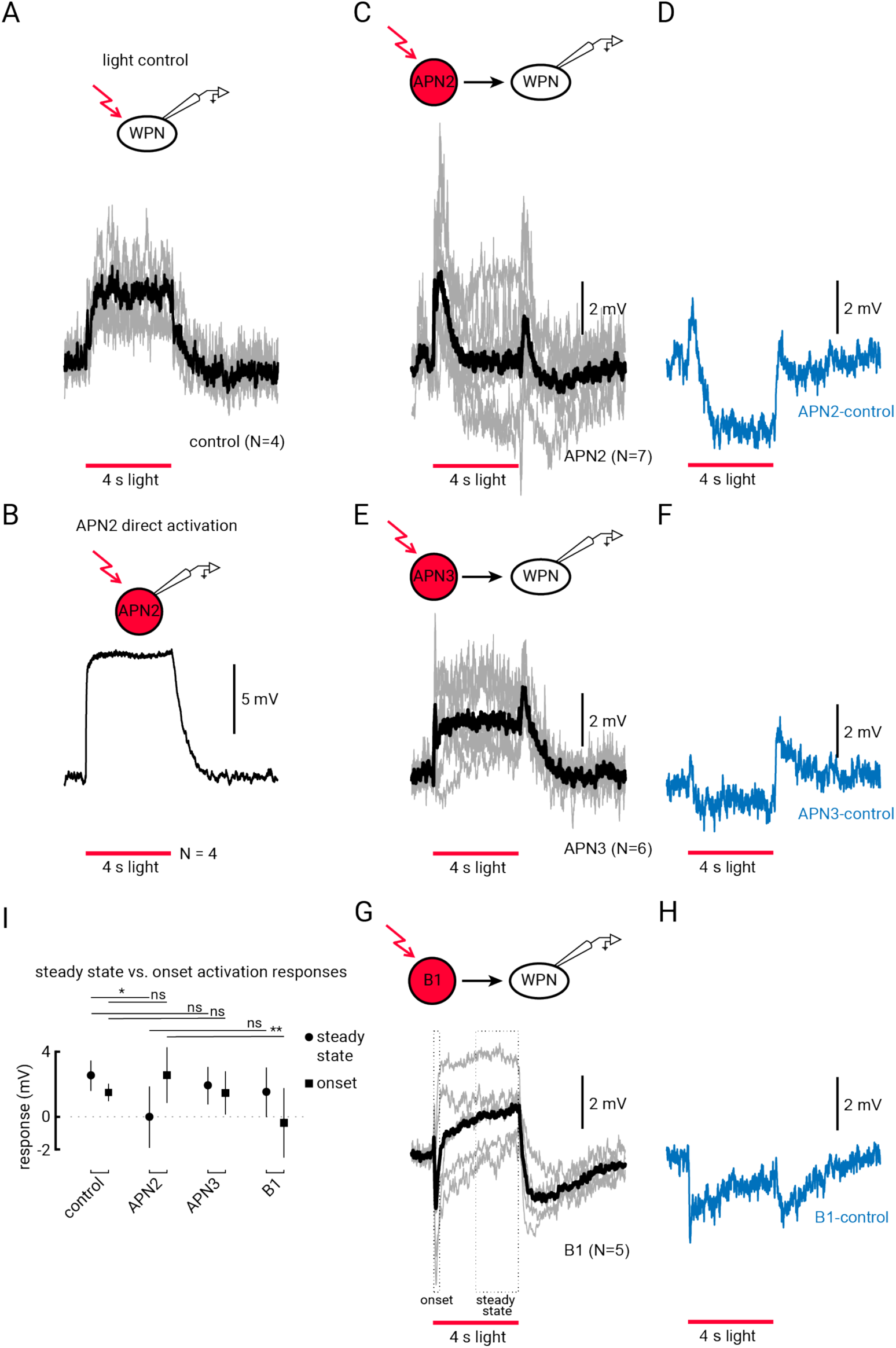
Response of WPNs to optogenetic activation of AMMC projection neurons. (A) Response of WPNs to light in control flies not expressing the light-activated channel. Individual fly averages are shown in grey, average across flies shown in black. Red bar indicates when the 4 s light stimulus is applied. (B) Average response of APN2 to direct optogenetic activation across N=4 flies (APN2>CsChrimson). (C) Average WPN response to light activation of APN2 (APN2>CsChrimson). Note the more transient response compared to control. (D) Difference between average control (A) and light-activation of APN2. (E) WPN response to light activation of APN3 (APN3>CsChrimson). (F) Difference between average control (A) and light activation of APN3 (E). (G) Average WPN response to light activation of B1 (B1>CsChrimson). Note initial hyperpolarization not seen in APN2 activation or control. (H) Difference between average control (A) and light activation of B1 (G). (I) Quantification of the light activation responses depicted in (A,C,E and G). Average steady state response (last 2 s during the light stimulus) are plotted as circles, and average onset response (peak response during 0.2 s near the beginning of the light stimulus) are plotted as squares. These regions are indicated above by the dotted squares in (G). Steady state response of APN2 is significantly different from control flies, but the onset response is not (two-sample t-test, P = 0.035 and P = 0.12). Steady state and onset responses of APN3 are not significantly different from control responses (two-sample t-test, P = 0.40 and P = 0.46). B1 onset response is opposite in sign and statistically significantly different from APN2 responses, but the steady-state responses are not different (two-sample t-test, P = 0.0083 and P = 0.17).

**Video S1. Confocal image stack of postsynaptic targets of JO-CE neurons using *trans-Tango***.

Green signal shows GFP signal from started line (JO-CE neurons), magenta labels postsynaptic targets, and neuropil is labeled in blue.

**Video S2. Confocal image stack of APN2s and their postsynaptic targets using *trans-Tango***.

Green signal shows GFP signal from started line (APN2s), magenta labels postsynaptic targets, and neuropil is labeled in blue.

**Video S3. Confocal image stack of APN3s and their postsynaptic targets using *trans-Tango***.

Green signal shows GFP signal from started line (APN3s), magenta labels postsynaptic targets, and neuropil is labeled in blue.

**Video S4. Confocal image stack of B1 neurons and their postsynaptic targets using *trans-Tango***.

Green signal shows GFP signal from starter line (B1 neurons), magenta labels postsynaptic targets, and neuropil is labeled in blue.

